# Physics-based, data-driven cell-scale membrane simulations with HMFF

**DOI:** 10.1101/2025.05.24.655915

**Authors:** Valentin J. Maurer, Marc Siggel, Rasmus K. Jensen, Julia Mahamid, Jan Kosinski, Weria Pezeshkian

## Abstract

Simulating entire cells represents the next frontier of computational biology. Achieving this goal requires methods that accurately describe cellular membranes across spatial and temporal scales. Although three-dimensional electron microscopy enables detailed membrane visualization, limitations on acquisition geometry, data quality, and field-of-view often result in fragmented membrane representations incompatible with simulations. To resolve this, here we introduce Helfrich Monte Carlo Flexible Fitting (HMFF), an approach that integrates experimental density data into physical simulations to determine membrane structure. Through the accompanying Mosaic software platform, we apply HMFF to influenza virus particles, *Mycoplasma pneumoniae* cells, and entire eukaryotic organelles. The resulting models enable multi-scale simulations spanning millions of lipids and proteins at experimentally determined positions, support quantitative morphological analysis, assess uncertainty in membrane localization, and reveal physical effects implicit in the data. Together, these capabilities establish a foundation for data-driven whole-cell simulations.

## I. MAIN

A cornerstone of modern biological research is deciphering cellular architecture and behavior, from individual molecules to the entire cellular landscape^1,2^. This requires an integrative approach that merges state-of-the-art experimental methods, such as cryogenic electron microscopy (cryo-EM)^3–5^, with cutting-edge computational advances, including multi-scale molecular simulations, from mesoscale to all-atom molecular dynamics (MD)^6,7^. These approaches operate in concert, overcoming individual limitations to bring us closer to building computationally tractable models resolving both structural and dynamic aspects of the cell^8–13^.

Within the scope of cellular architecture, biomembranes represent a particularly complex and essential component that requires specialized modeling strategies. Biomembranes play a vital role in many cellular processes such as signaling, transport, and compartmentalization across all domains of life. They form complex and highly curved structures, spanning from nanometer to micrometer scale^14–16^. Their constant turnover and reshaping occur over time scales ranging from nanoseconds to minutes. Resolving membrane architecture, however, is a challenging task, both experimentally and computationally^6,14,17,18^. While cryo-EM provides valuable structural details, it suffers from resolution limitations and provides only static snapshots of inherently dynamic systems^3,19–23^. On the other hand, molecular modeling faces challenges such as insufficient information about membrane composition, timescale limitations, and system parameter uncertainties^6,7,18^. A promising approach to accurately determine membrane structure is to develop an integrative modeling approach, which combines physics-based modeling techniques with experimental data as constraints to infer structural details that remain hidden from experiments alone.

To achieve this, here we introduce Helfrich Monte Carlo Flexible Fitting (HMFF), which integrates volumetric density data directly into dynamically triangulated surface (DTS) mesoscale simulations of biomembranes^7,24,25^. HMFF extends the physical energies underlying DTS simulations, most notably the Helfrich bending energy^26^, with a data-derived term that transfers the principle of Molecular Dynamics Flexible Fitting^27–30^ from atomic structures to mesoscale membranes and includes a membrane-specific coherence regularization for stability. In the combined energy landscape, experimental data guides membranes toward observed densities while physical energies regularize and interpolate their shape. DTS samples this landscape through Monte Carlo moves, producing a thermodynamic ensemble of configurations rather than a single minimum-energy shape, capturing fluctuations relevant to emergent, dynamic membrane behavior (in the thermodynamic sense) such as protein reorganization^31^ and curvature-mediated membrane folding^32^. To enable HMFF on experimental data, we developed Mosaic, a software platform comprising the necessary methods delivered through a graphical user interface. Mosaic integrates established tools, including MemBrain-seg for segmentation^19^, FreeDTS for simulation^24^, PyTME for constrained template matching^33^, and TS2CG for backmapping to coarse-grained representations^34,35^, with algorithms for mesh completion, protein projection, and morphometric analysis.

We demonstrate our approach across biological systems, length scales, and experimental methods. We validate HMFF on synthetic reference systems with known ground truth and on real cryo-ET data through global and local ablation experiments. We simulate membranes of partially imaged *Mycoplasma pneumoniae* cells and filamentous influenza A virus (IAV) from cryo-ET data, and organelles throughout an entire eukaryotic cell imaged by focused ion beam scanning electron microscopy (FIB-SEM). We refine these mesoscale models into molecular-level representations^34,35^, facilitating simulations spanning from individual membrane proteins to systems with millions of lipids. Together, these applications bridge static experimental observation and dynamic physical simulation, enabling simulations of complex membrane systems with shapes representative of those observed in nature. This represents a concrete step toward whole-cell modeling.

## II. RESULTS

### A. Integrating experiment and simulation using HMFF

We developed HMFF to create models of biomembranes that are consistent with experimental observation and physical membrane properties. Our approach builds on DTS simulations^24,25^, in which membranes are modeled as triangular meshes whose configurations are governed by the Helfrich model^26^. The associated bending energy penalizes deviations from preferred curvature, favoring smooth shapes, and can be extended with biologically relevant terms for volume, membrane protein inclusions, or interactions with external systems such as cytoskeleton or extracellular matrix^24^. To this physical model, HMFF adds an energy term^27–30^ that guides the simulation towards membrane densities in experimental data. In the combined energy landscape, physical energies regularize and interpolate membrane shape while experimental density provides structural guidance.

To illustrate the concept, we simulated a planar membrane embedded in a Gaussian density mimicking a membrane signal (Fig. 1a, Mov. M1). The HMFF energy drove the membrane toward the high-density region, curving it transiently against its bending resistance before it equilibrated in its preferred flat configuration. The coupling parameter *ξ* governs the balance between data and physical energies (Fig. 1b). Increasing *ξ* led to faster convergence toward high-density regions, while values approaching zero failed to displace the membrane from its initial position. In the limits, *ξ* → ∞ yields a purely data-driven simulation and *ξ* → 0 reduces to the physical model.

**Fig. 1.**
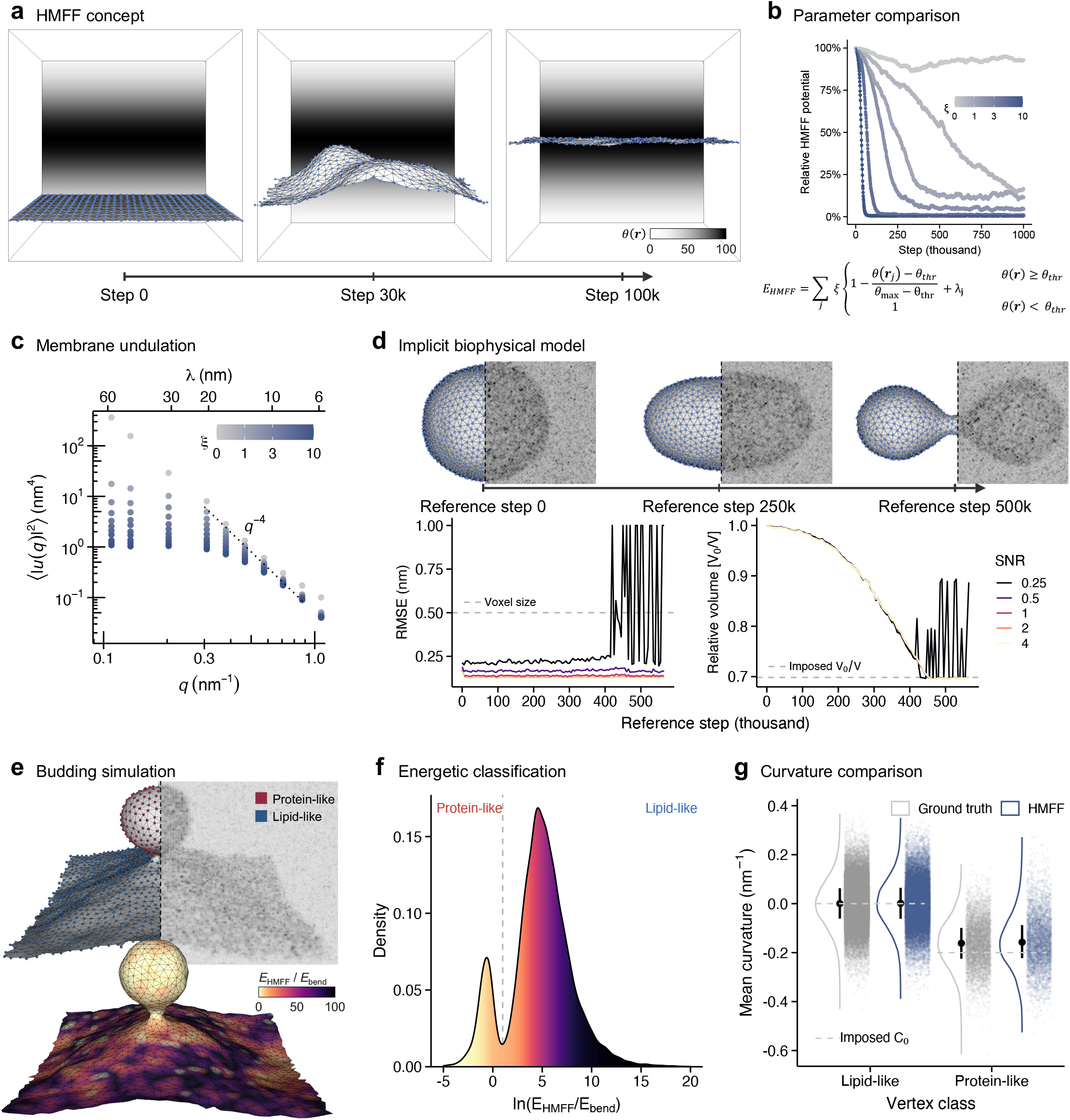
HMFF concept and validation on synthetic systems. **a**, Planar membrane simulation with periodic boundary conditions embedded in a Gaussian density *θ*(**r**) projected to the back plane of the box. **b**, HMFF potential along the simulation trajectory for different coupling strengths *ξ*, min-max scaled per *ξ*. **c**, Equilibrium undulation spectrum ⟨|*u*(*q*)| ^2^⟩ of the membrane in **a** for the indicated *ξ*. Average of 4.5 million simulation steps. Dotted line: theoretical *q*^−4^ scaling. **d**, Membrane simulation under volume and curvature coupling. Corresponding simulated density at SNR 0.5 shown as maximum intensity projection (MIP, top). HMFF simulations started from a sphere and were run against simulated densities of each reference step at the indicated SNR. RMSE to the reference (left) and relative volume (right), averaged over 50 thousand simulation steps and three replicates; dashed lines mark the voxel size of the simulated density and the imposed volume ratio. RMSE values beyond 1 nm were truncated. **e**, Budding driven by protein inclusions with spontaneous curvature *C*_0_. Reference DTS simulation, with vertices colored by vertex class, and MIP of the corresponding density at SNR 1.0 (top); HMFF simulation starting from a membrane bud against a simulated density of the reference at SNR 1.0 (bottom). Color scale indicates quantiles of *E*_HMFF_*/E*_bend_. **f**, Distribution of *E*_HMFF_*/E*_bend_ across the HMFF mesh in **e**, over 15 thousand simulation steps, partitioning vertices into protein-like and lipid-like classes. **g**, Mean curvature per vertex class for reference and HMFF simulation. Dashed lines indicate the imposed spontaneous curvature *C*_0_. Data shown as median and interquartile range (IQR).

To understand how *ξ* shapes membrane behavior, we examined the undulation spectra of the HMFF-driven flat membrane system (Fig. 1c). The Helfrich model predicts a characteristic undulation spectrum, which has been confirmed by molecular simulations^36^ and experiments^37^. Our spectra matched the theoretical prediction at short wavelengths and deviated above a crossover wavelength, where the density coupling outweighs bending and biases global shape. Increasing *ξ* shifted this crossover to shorter wavelengths. This analysis demonstrated that HMFF samples a thermodynamic ensemble of physically valid configurations rather than collapsing to a single optimization minimum, retaining membrane fluctuations below a crossover wavelength dictated by *ξ*.

HMFF acts as an implicit model of physical effects embedded in the experimental data, capturing factors that would otherwise require explicit parametrization. To demonstrate this, we simulated a spherical membrane subject to osmotic pressure and bilayer asymmetry, modeled by volume, area, and global curvature coupling (Fig. 1d, top)^24^. For each frame of this reference trajectory, we generated a simulated density (Fig. S1a) and ran an HMFF simulation against it (an HMFF simulation refers to a DTS simulation with the HMFF energy), starting from the initial sphere and without explicit volume coupling. HMFF recovered both the reference geometry and the volume ratio imposed by volume coupling in the reference simulation across all frames from the density alone, down to a signal-to-noise ratio (SNR) of 0.5 (Fig. 1d, bottom). This similarly applied to another system (Fig. S1b)^34^. Recovery quality also depended on *ξ*, with higher *ξ* helping overcome energy barriers but at the risk of biasing the mesh toward noise in the data. To mitigate this, the HMFF energy includes a coherence regularization term *λ*, which forces adjacent parts of the mesh to occupy regions of similar density. This regularization substantially reduced Root Mean Square Error (RMSE) standard deviation at higher *ξ*, broadening the range over which recovery is stable (Fig. S1c).

Building on this implicit modeling capability, we next asked whether we could visualize where these unknown physical effects act on the membrane. We simulated a planar membrane containing curvature-inducing proteins^24^, which drive membrane budding (Fig. 1e, top). We then held the resulting bud in place against its simulated density, without modeling the proteins. We reasoned that relating the HMFF energy to the physical energies governing the simulation would reveal where the data drives the membrane against its physical preferences, highlighting regions where an effect absent from the physical model acts and the data must compensate. Since the omitted proteins acted through curvature, we examined the per-vertex ratio *E*_HMFF_*/E*_bend_ (Fig. 1e, bottom), which quantifies where the data acts against membrane bending preference. The distribution of *E*_HMFF_*/E*_bend_ was bimodal, partitioning vertices into protein-like and lipid-like classes (Fig. 1f), with low and high ratios, respectively. The mean curvature of each class matched the corresponding reference distribution (Fig. 1g; KS statistic *D* = 0.01 for lipid and 0.046 for protein), confirming that the energetic imbalance identifies protein inclusion locations from density data. Demonstrated here for membrane proteins, this generalizes to other effects on membrane shape, providing a means to localize their action from the data.

### B. From experimental data to HMFF simulation with Mosaic

While HMFF performs data-driven membrane simulations, applying it to experimental data requires specialized methods from setup to analysis. To realize this workflow, we developed Mosaic, a software platform accessible through a graphical user interface (GUI, Fig. 2a). We illustrate it on cryo-ET data of *M. pneumoniae* cells^38,39^.

**Fig. 2.**
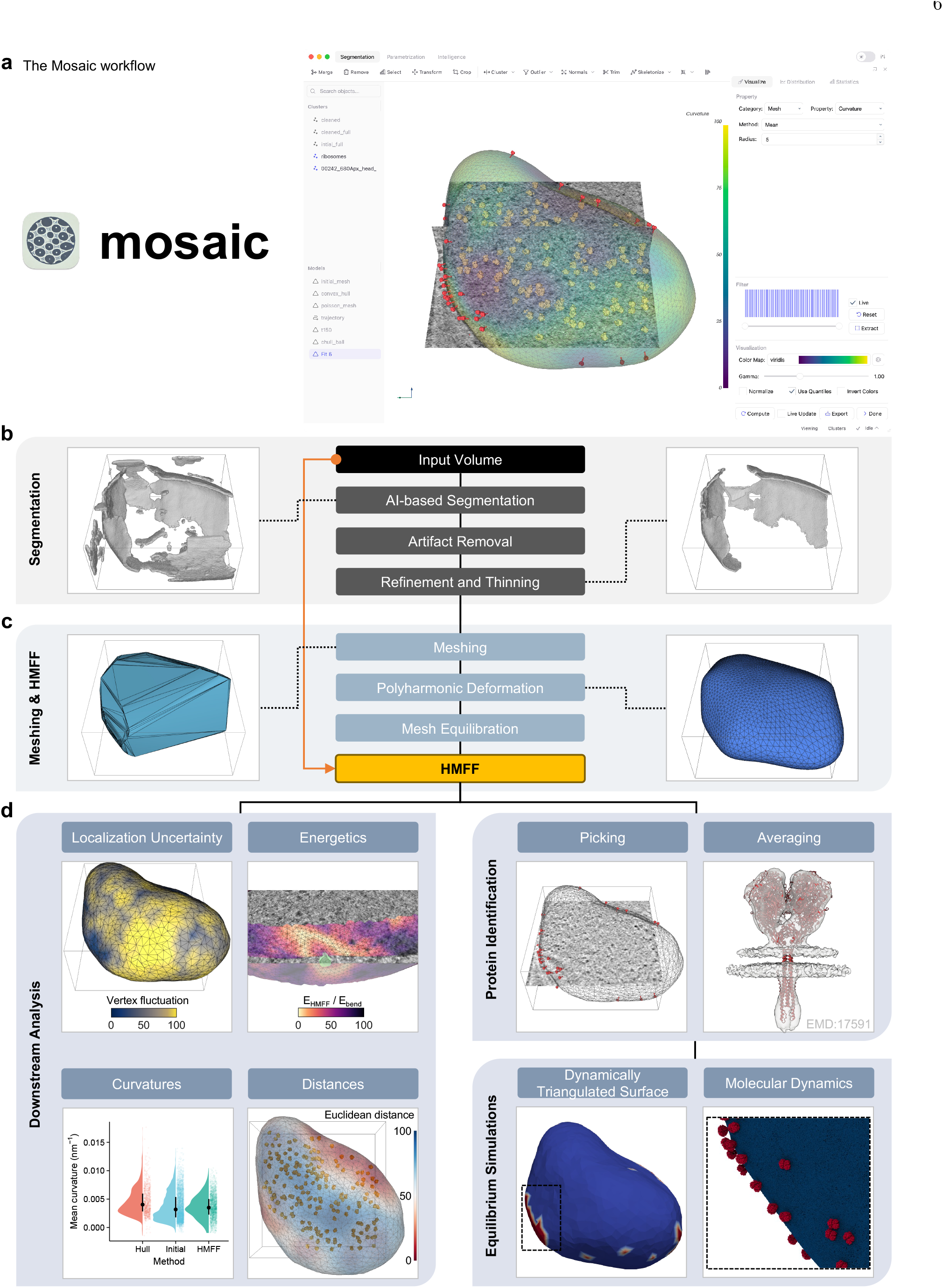
Mosaic workflow illustrated on *M. pneumoniae* cryo-ET data. **a**, Mosaic user interface. **b**, Interactive refinement of membrane segmentations. **c**, Creating meshes for HMFF simulation. **d**, Downstream applications. Left, per-vertex maps colored by localization uncertainty (vertex fluctuation across the ensemble) and energetics (*E*_HMFF_*/E*_bend_). Below, distributions of mean curvature (median ± IQR) and of Euclidean distance to ribosomes, each computed for the convex hull of the segmentation (Hull), the polyharmonic-deformed mesh (Initial), and the HMFF mesh. Color encodes quantiles throughout. Right, Nap complexes picked using the HMFF mesh. The picks can be subtomogram averaged (EMD:17591 shown for illustration), combined with the HMFF mesh for equilibrium DTS simulations, and jointly backmapped to a coarse-grained Martini representation for MD simulations. Tomogram was bandpass-filtered for visualization.

Mosaic integrates deep-learning-based membrane segmentation^19^ with interactive cleanup tools (Fig. 2b, S2a-b). Using these tools, we removed mis-segmented regions in the cytosol and at tomogram boundaries and thinned the membrane to single-voxel thickness for meshing (Fig. S2c, Mov. M2). The cell extended beyond the tomogram boundaries, leaving only opposite sections of the membrane visible. This caused standard meshing approaches such as Poisson reconstruction^40–42^ to fail (Fig. S2f). Thus, we developed a mesh completion procedure based on polyharmonic deformation that produces closed meshes while preserving smoothness and local curvature of the observed membrane (Fig. 2c, S10). The resulting mesh was 43% (0.091 µm^3^) larger in volume than the convex hull of the segmentation and, unlike the convex hull, encapsulated all ribosomes, indicating that the completion captures the cell geometry more accurately. DTS simulation imposes mesh quality criteria such as bounded edge lengths and the absence of self-intersections. While our meshes already met most of these, we devised a two-step equilibration combining remeshing and hybrid MC to regularize edge lengths (Fig. S2e). We then used HMFF to simulate the membrane against the tomogram. Our initial simulation assumed zero exterior density, creating an energetic imbalance that caused the mesh to adhere to the tomogram boundary (Mov. M4). We resolved this through a dedicated padding procedure (Fig. S2g), yielding a mesh that captured the experimentally observed membrane while interpolating beyond the tomogram boundaries based on physical membrane properties (Fig. 2d, S3, Mov. M3).

To validate the application of HMFF on experimental data, we performed two ablation experiments against manually annotated *M. pneumoniae* cells^20^ (Fig. S4a). In a global ablation, we progressively replaced experimental density with random values from the tomogram, testing HMFF’s tolerance to continuous degradation of the data signal. HMFF maintained accuracy up to approximately 90% randomization (Fig. S4b). In a local ablation, we masked rectangular regions, removing the membrane signal and forcing reliance on physical interpolation. HMFF maintained accuracy for mask sizes up to 40 voxels (RMSE ≤ 2 nm) and degraded at 80 voxels (109 nm), with the steepest deterioration in convex regions (Fig. S4c).

Building on this validation, we used the HMFF mesh of the *M. pneumoniae* cell for downstream simulation and analysis (Fig. 2d). HMFF produces a thermodynamic ensemble of valid membrane configurations, whose fluctuations can be used to quantify uncertainty in membrane localization (Fig. 2d, S4d). The fluctuations are governed by the energies underlying the simulation, which can be decomposed to reveal where the data deviates from the physical model. Since the physical model here consisted only of the bending energy, we investigated the ratio *E*_HMFF_*/E*_bend_. We found a localized region of elevated ratio that coincided with a pronounced membrane deformation and an external particle with apparent polyhedral geometry (Fig. S4e), demonstrating how such energy decomposition can reveal physical effects implicit in the experimental data.

The HMFF mesh also supports classical geometric analyses. The mean curvature distribution changed during HMFF simulation (Fig. 2d), illustrating the interplay between data and physical energies – since the bending-energy minimum for a closed surface corresponds to a uniformly curved sphere, the variance in the resulting distribution reflects the contribution of the HMFF energy. Distance-based analyses further quantify membrane association of cellular features such as ribosome localization (Fig. 2d). The HMFF mesh can also serve as a starting point for multi-scale simulation of membrane-protein systems. As an example, we mapped positions of Nap, a transmembrane protein complex essential for infectivity^43^ by applying constrained template matching^33,44^, which uses the HMFF mesh to restrict particle picks to physically plausible configurations on the membrane. Consistent with previous reports^43^, Nap concentrated around the specialized attachment organelle. Combining the HMFF mesh with Nap picks, we built an *in silico* model representing each Nap complex as a membrane inclusion. Given a parametrization of how Nap influences local membrane properties, this model can probe emergent behavior such as curvature-mediated clustering and sorting of proteins^32^. The same model can be refined to higher resolution, where interactions are modeled through molecular force fields rather than the effective parameters of the DTS model. To demonstrate this, we backmapped it to a coarse-grained Martini representation^34,35^, yielding a complete cell membrane with 44 Nap complexes (approximately 50 million particles) that we briefly equilibrated by MD^45,46^. This places the workflow across multiple resolutions, from the DTS mesoscale to near-atomistic representations.

Together, these results demonstrate that Mosaic enables the application of HMFF to experimental cryo-ET data, quantitative analysis of the resulting membrane ensembles, and downstream multi-scale simulations.

### C. Simulating filamentous influenza A virus-like particles

Influenza A virus (IAV) is pleomorphic, ranging from spherical to filamentous phenotypes. Computational modeling has so far concentrated on spherical particles^47–49^, leaving filamentous IAV unmodeled despite their clinical significance^50^. Beyond being consider-ably larger than their spherical counterparts, filamentous IAV carry tightly packed hemagglutinin (HA) and neuraminidase (NA) glycoproteins on the surface and a dense matrix protein layer (M1) lining the inner membrane. Building a model that reflects this native architecture, rather than an idealized geometry, requires identifying the membrane and its proteins directly from the data, a task complicated by their dense and heterogeneous packing.

To build a filamentous IAV model, we used publicly available cryo-ET data of filamentous virus-like particles (VLPs)^51^. Fig. 3a illustrates the workflow from tomographic data to HMFF mesh, with intermediate steps in Fig. S5a-d. The membrane and M1 protein layers lie in close proximity, and segmentation failed to differentiate them. Instead, the segmentation primarily captured the M1 layer, presumably because its elongated high-density appearance mimics a membrane (Fig. 3a, Mov. M5). This propagated to the initial mesh, which was positioned in the mid-plane of the M1 layer rather than on the membrane.

**Fig. 3.**
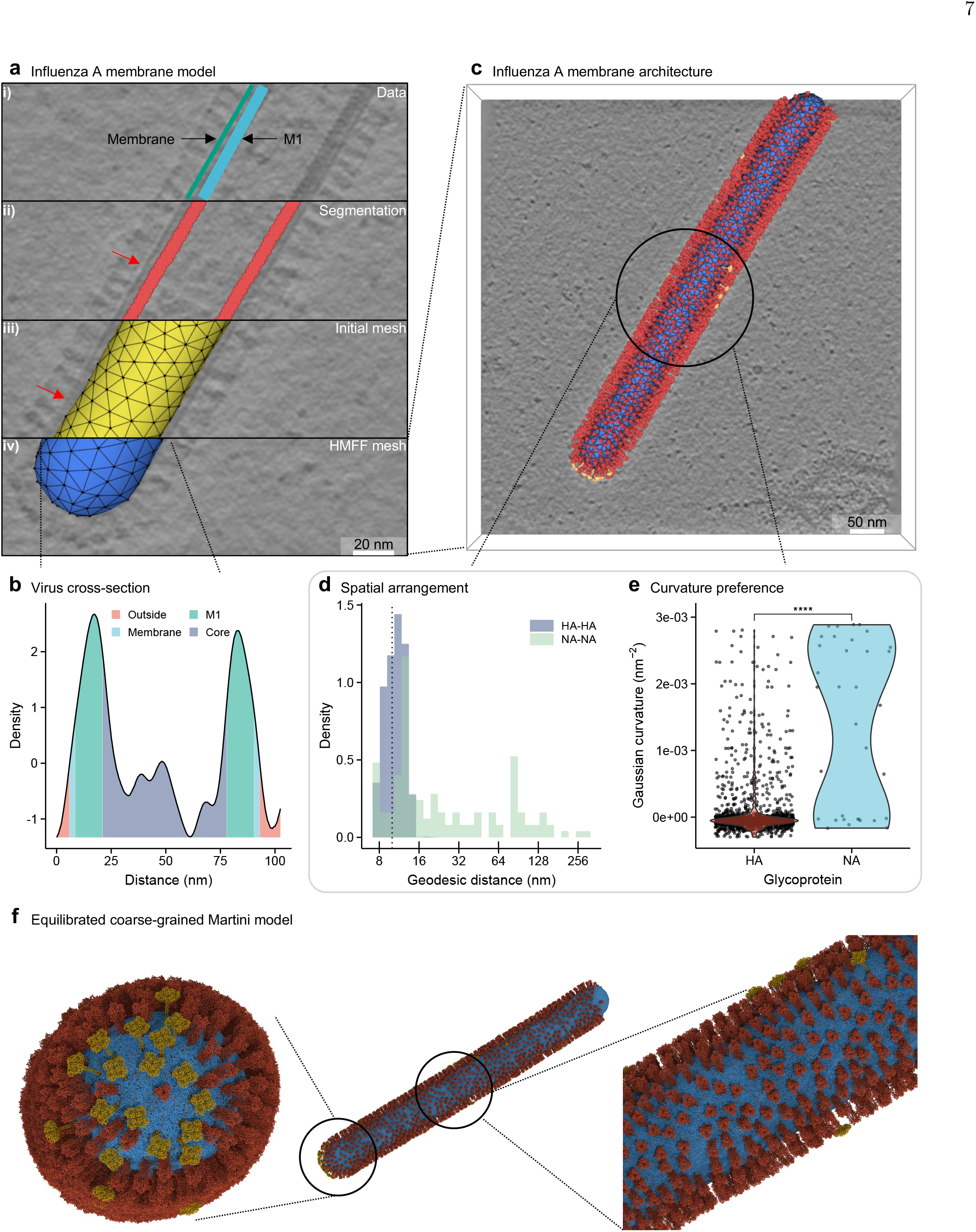
HMFF simulation and analysis of filamentous IAV. **a**, (i) tomogram (EMD:11075) with membrane and M1 protein layer annotated; (ii) segmentation (red) primarily capturing the M1 layer, deviation from membrane is indicated by red arrows; (iii) initial mesh (yellow) sitting on the M1 mid-plane; (iv) HMFF mesh (blue) obtained with volume coupling. Tomogram was deconvolved using mtffilter^55^. **b**, Density profile of the VLP cross-section indicated by the dotted line in **a**. Core refers to the content of the VLP not covered by other zones. **c**, Membrane architecture of the VLP. HMFF mesh (blue) with backmapped HA (red) and NA (orange) picks. **d**, Distribution of HA-HA and NA-NA geodesic distances to four nearest neighbors. **e**, Gaussian curvature at HA and NA locations (one-tailed Mann-Whitney-U-Test; \*\*\*\**p <* 0.0001). Data shown as median ± 1.5 IQR. **f**, Equilibrated coarse-grain model combining picks and HMFF mesh. Shown is a view of the filament cap (left), side view (middle), and a magnified section of the side view (right). Beads colored by component: red (HA), yellow (NA), blue (membrane). One filament end lacks proteins due to tomogram boundaries.

In our initial simulation, the mesh collapsed onto the M1 layer, which is denser than the membrane in cross-section (Fig. 3b) and therefore exhibits a greater attraction. Because the mesh began at the M1 mid-plane interior to the membrane, it underestimated the true volume of the VLP. Thus, we encoded this observation in the simulation through volume coupling, modeling the mesh as having a higher osmotic pressure than its surroundings. With this enrichment of the physical model, the simulation converged onto the membrane (Fig. 3a iv, Mov. M6). In the preceding sections, we treated the mesh edge length as a fixed parameter. It has a more fundamental role, however, bounding the finest length scales the simulation can resolve, much like a Nyquist limit. Reducing it from 11 nm to 7 nm, along with adding *λ* regularization, reproduced the volume-coupled mesh without an explicit volume term (Fig. S6a, RMSE 1.71 nm), indicating that the information distinguishing membrane from M1 is present in the density, but at a different length scale than initially assumed.

To assess the accuracy of the HMFF mesh, we used HA picks from constrained template matching (Fig. 3c, S5e-i) as a positional reference. We measured the distance and angular deviation of the picks to the mesh, and compared the distributions to those of the segmentation and a cylindrical approximation of the complete VLP. We restricted the comparison to HA to avoid penalizing the cylinder at the spherical caps where it cannot fit (Fig. 3c). Distances from HA picks to the HMFF mesh most closely matched the expected value from the atomic structure (Fig. S6b). 90% of HA particles deviated by less than 20° from the surface normal (Fig. S6c), consistent with near-perpendicular insertion, and the distribution was narrower than for the segmentation. Together, these results indicated that the HMFF mesh accurately recovers both the position and curvature of the membrane, providing a better starting representation for downstream applications such as template matching and subtomogram averaging.

Building on this mesh accuracy, we projected HA and NA picks onto the HMFF mesh to characterize glycoprotein organization on the viral envelope (Fig. S5e-i, S6d). HA densely populated the cylindrical shaft with HA-HA spacing narrowly distributed around 10 nm (Fig. 3c-d), consistent with previous cryo-ET analyses^50,52,53^. NA showed a bimodal pattern, with one population exclusively occupying the spherical caps and a second along the shaft (Fig. 3c). This bimodality was reflected in both the broader, more variable NA-NA spacing (Fig. 3d) and the NA curvature distribution (Fig. 3e), hinting at additional drivers of localization beyond curvature alone. Combining the HMFF mesh with the projected protein picks, we assembled and equilibrated a coarse-grained Martini representation of the filamentous IAV envelope (10 million particles, Fig. 3f, Mov. M7), comparable in size to recent large-scale systems^47,54^, providing a foundation for future molecular simulation.

### D. Scaling to whole eukaryotic cells

Eukaryotic cells occupy a markedly different regime than the systems considered so far. They contain diverse organelles, volumes orders of magnitude larger and more complex morphologies, from branched mitochondria to tubular endoplasmic reticulum. To examine whether our methodology can describe this complexity and scale, we applied it to FIB-SEM data of an entire HeLa cell^22^ measuring more than 30 µm in diameter.

We processed HeLa cell segmentation labels from the OpenOrganelle database, spanning mitochondria, Golgi apparatus, nucleus, endoplasmic reticulum, and plasma membrane (Fig. 4a-c, Mov. M10). The initial segmentations contained topological issues incompatible with simulation, including gaps, incorrectly merged objects, and artificial invaginations (Fig. S7a-i). Using Mosaic’s interactive tools, we resolved these issues, producing 237 mitochondrial meshes that preserved the underlying morphology and were suitable for subsequent simulation (Fig. 4c).

**Fig. 4.**
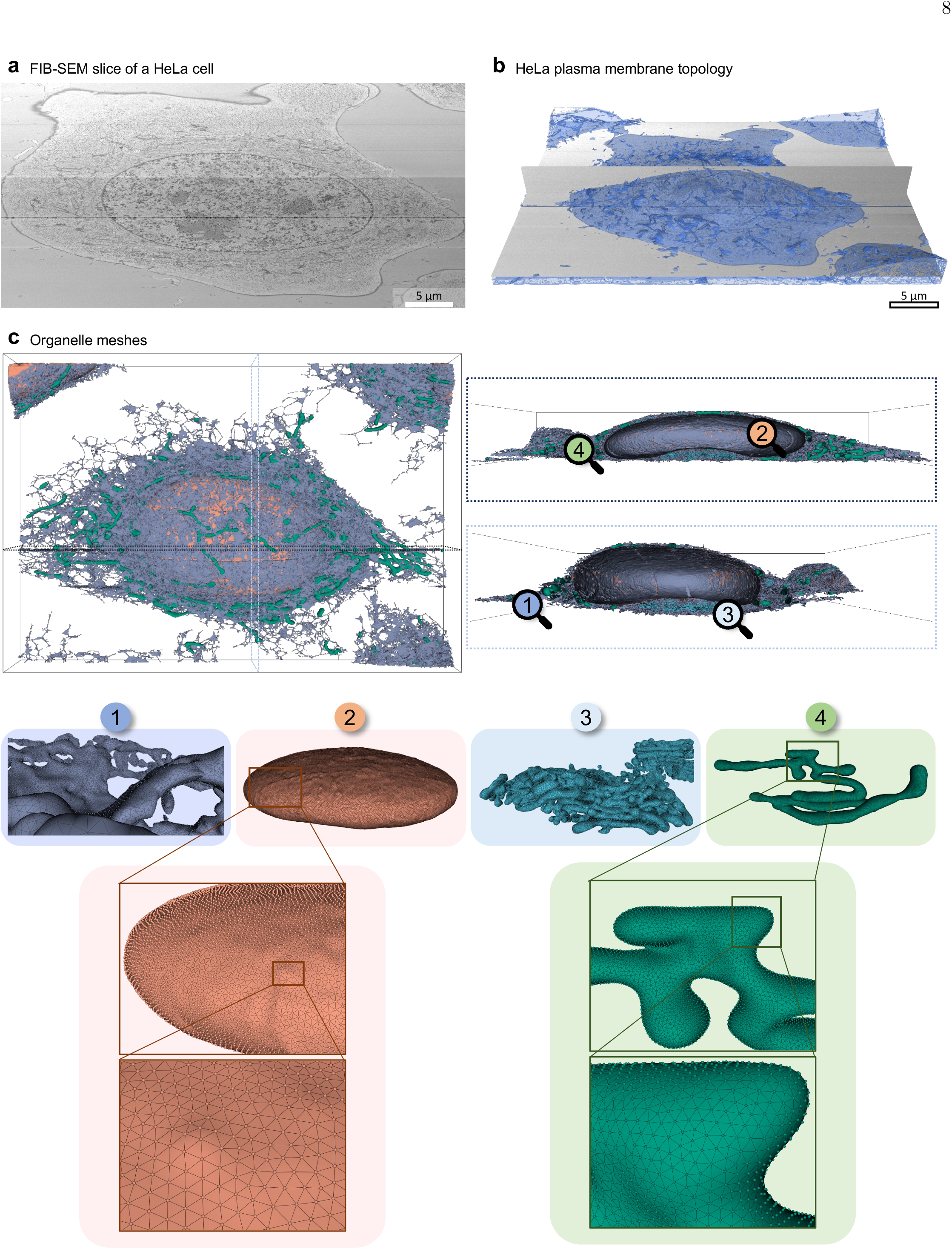
Triangulation of an entire HeLa cell. **a**, Top-view of FIB-SEM data (OpenOrganelle:jrc hela-2). **b**, Side-view of FIB-SEM data with plasma membrane mesh overlaid (blue). **c**, 3D rendering of triangulated cellular components with corresponding perpendicular slices shown on the right: (1) endoplasmic reticulum (ER) with complex tubular network, (2) nucleus, (3) Golgi apparatus, and (4) mitochondria with branched morphology.

At equilibrium, the HMFF mesh closely followed the outer mitochondrial membrane contours, particularly in regions of high curvature (Fig. 5a-b, Mov. M8, M9). During simulation, the HMFF energy, surface area, and volume decreased in tandem, consistent with the mesh contracting onto the dense membrane without distorting its shape (Fig. S8a-b). In previous sections, we showed that different simulations can produce comparable outcomes. To test this systematically, we performed a grid scan in the bending rigidity *κ* and the coupling strength *ξ*, which revealed a corridor of distinct parameter combinations yielding meshes closely matching that in Fig. 5a by RMSE (Fig. 5c).

**Fig. 5.**
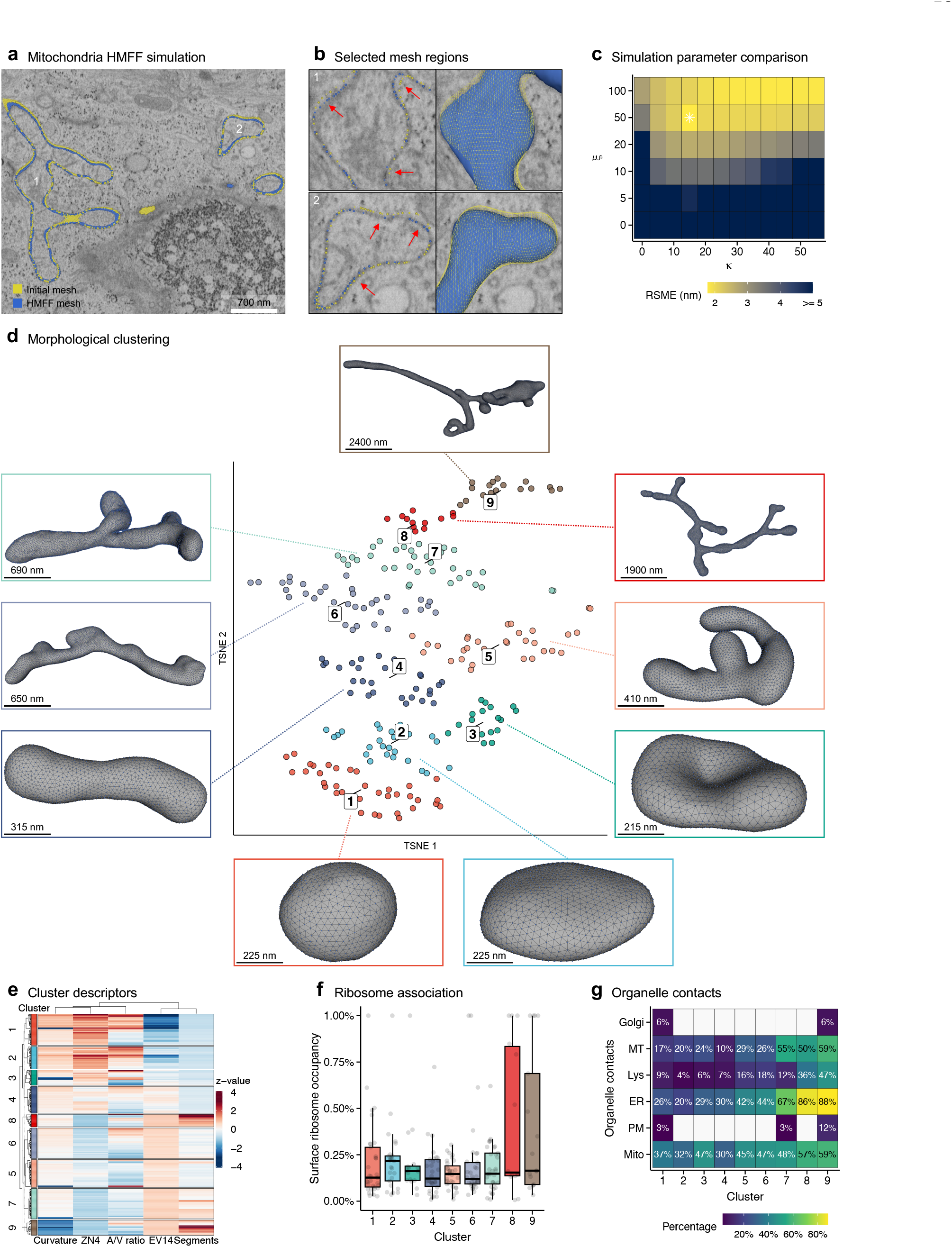
HeLa mitochondria simulation, classification, and characterization. **a**, Comparison of initial and HMFF mesh on a FIB-SEM slice. **b**, Magnified regions from (**a**); red arrows highlight where the HMFF mesh follows the membrane more accurately than the initial mesh. **c**, RMSE between the mesh in (**a**, asterisk) and meshes from a grid of bending rigidity (*κ*) and coupling strength (*ξ*), averaged over 10 thousand simulation steps. **d**, t-SNE embedding of mitochondrial descriptors, colored by Louvain cluster. **e**, Heatmap of the most discriminative descriptors by between-/within-cluster variance: area/volume ratio, 14th eigenvalue (EV14), segment counts, 4th Zernike moment (ZN4), Gaussian curvature. **f**, Mitochondria surface area in contact with ribosomes, per cluster (median ± 1.5 IQR). **g**, Percentage of mitochondria in contact with mitochondria (Mito), lysosomes (Lys), plasma membrane (PM), endoplasmic reticulum (ER), and microtubules (MT), per cluster. Values beyond 1% were truncated.

We next turned to the mitochondrial population as a whole, asking whether the morphological heterogeneity characteristic of mitochondria^56^ would resolve into distinct subpopulations across the 237 meshes reconstructed from the HeLa cell. From each mesh, we computed complementary descriptors capturing topological, geometric, and scale-dependent aspects of morphology (see methods IV N). Applying t-Distributed Stochastic Neighbor Embedding (t-SNE) dimensionality reduction followed by Louvain community detection revealed nine clusters with distinct structural signatures (Fig. 5d–e, S8c, cluster sizes reported in Tab. S1). The clusters ranged from compact, ovoid mitochondria with minimal branching (clusters 1–3) to extensively branched, network-like structures (cluster 9), with intermediate forms including elongated tubular (4), Y-branched (5), multi-junction (6), sheet-like (7), and asymmetric, dilated (8) morphologies.

We examined whether these clusters reflect functional differences. Using ribosome segmentations from the original dataset^22^, we found that ribosome-mitochondria association did not differ significantly across clusters (two-sided Mann-Whitney-U-Test, all *p >* 0.05 using Bonferroni correction; Fig. 5f), consistent with the general requirement for ribosome association in mitochondrial biogenesis. Organelle contact patterns, in contrast, varied across clusters (Fig. 5g). Inter-mitochondrial contacts were observed in clusters 5-9; ER-mitochondria contacts spanned the same range and peaked in clusters 8 and 9 (86% and 88% of mitochondria, respectively); microtubule associations exceeded 50% in clusters 7-9; and lysosome contacts were highest in clusters 8 and 9. Several of these associations were enriched in the most topologically complex clusters, suggesting functional specialization linked to morphological complexity.

Together, we demonstrated that our approach generalizes to FIB-SEM data of entire eukaryotic cells, producing mesh representations that both support simulation of micrometer-scale organelle membranes and enable morphological characterization at the population scale, extending the analyses of earlier sections.

## III. DISCUSSION

Membrane structure has remained difficult to determine from experiment or physical simulation alone, as each resolves only part of it. HMFF overcomes this limitation by integrating the two. To take HMFF from theory to experimental practice, we developed Mosaic, a software platform that streamlines the setup and downstream analysis of HMFF simulations. Together, HMFF and Mosaic now let us simulate and characterize biomembranes across scales, from nanometer features to micrometer-scale cellular compartments.

Where segmentation and morphometric analysis describe the experimentally observed surface^19,20,41^, HMFF produces a thermodynamic ensemble that captures the many microstates a membrane explores in reality, which a single configuration cannot, regularizing shape and interpolating regions unobserved in the data; where large-scale simulations rely on idealized geometries and protein distributions^1,34,54,57,58^, our approach yields shapes representative of those in nature with proteins at experimentally determined positions, suitable for simulation of membrane–protein systems from the mesoscale to near-atomistic resolution. Such models do not merely reproduce the data but go beyond it, separating the IAV membrane from the adjacent M1 layer to describe the virus more faithfully than alternative representations or completing partially imaged *M. pneumoniae* cells.

Our models of membrane structure generate testable biological hypotheses. Decomposing the *M. pneumoniae* simulation into its energetic constituents revealed an external particle co-localizing with a pronounced membrane indentation, a physical effect implicit in the data but absent from the physical model. Both its polyhedral geometry, reminiscent of cryo-ET-resolved lipoprotein particles^59^, and the indentation are consistent with a lipoprotein engaging the host-lipid-scavenging machinery on which *M. pneumoniae* depends^60,61^. Such observations can be interrogated through the ensemble, which reflects the underlying energy landscape and provides a measure of localization uncertainty. While NA clustered at the caps of the IAV envelope^52,62^, its global distribution departed from strict polarity, consistent with fluorescence microscopy observations^50^. This organization may reflect the stochastic nature of membrane mechanics rather than additional drivers, a distinction resolvable by parametrizing HA and NA as inclusions and testing whether simulation produces an ensemble recapitulating the observed distribution. The HeLa mitochondrial population defines an empirical shape space, analogous to the theoretical shape spaces that membrane physics predicts under defined conditions^26,63^. By this analogy, each morphology may arise from a distinct energetic regime, a physical signature that simulation can probe by searching for parameters that reproduce the observed shape.

We see three key limitations of our approach. First, in regions where data is genuinely absent, such as membranes perpendicular to the electron beam in cryo-ET or areas beyond the field of view, membrane structure is determined by the physical model alone. The accuracy of this interpolation depends on simulation parameters that interact non-trivially through the set of accessible mesh configurations, so that the mapping from parameters to shape is many-to-one. The IAV simulation illustrated this, as reducing the mesh edge length produced a shape that, at the scale of membrane thickness, was indistinguishable from the one obtained with explicit volume coupling. The edge length sets the length-scale range accessible to the mesh, here phenomenologically reproducing the effect of an additional physical energy. While Mosaic provides utilities for screening and analyzing parameter sets, the degeneracy in DTS parameters currently precludes analytical procedures for translating HMFF’s implicit parametrization into explicit biophysical parameters. As a consequence, removing the HMFF energy causes the membrane to relax from the observed shape, unless the physical model is sufficient to support it. Second, DTS simulations have a practical size limit. While CPU parallelization alleviates this, whole-eukaryotic-cell simulation at functional membrane resolution remains beyond current reach. Third, and more broadly, topology changes such as fusion or fission events present a challenge inherent to DTS simulations. These involve large energy barriers and dedicated biological catalysts, and whether minimizing free energy alone produces biologically realistic topologies remains an open question. In principle, the HMFF energy could be repurposed to inform simulations through such events, though this lies beyond current capabilities.

Several future directions could strengthen the approach. The data guiding the simulation could be improved by denoising to strengthen the membrane signal, weighting by local reliability, or incorporating orthogonal information such as protein localizations. Approaches to calibrate physical parameters such as protein curvature preference, whether from experiment or from the equilibrium structure of the simulation itself, would enable the study of realistic protein dynamics. Further afield, serial lift-out can image near-complete organisms as a series of slices^64,65^, which HMFF could render continuous by anchoring the observed regions and interpolating across the gaps left by material loss during extraction. Coupling the two would extend the approach from organelles to organisms and make questions currently out of reach tractable, such as how membrane remodeling at the cellular scale during viral infection propagates across the tissue.

In summary, HMFF and Mosaic close a long-standing gap between what can be imaged and what can be simulated for biomembranes, advancing the field toward the broader goal of creating foundational models of whole cells.

## IV. METHODS

### A. Dynamically triangulated surface simulations

At a scale much larger than its thickness, a membrane can be modeled as a two-dimensional surface in three-dimensional space. By the Fundamental Theorem of Surface Theory, given the metric and the curvature tensor, the surface can be reconstructed uniquely up to rigid motions, i.e., translations and rotations. Since the membrane surface is related to the number of biomolecules and is an extensive variable, the bending energy of the membrane, up to second order in curvature, can be well described by the Helfrich Hamiltonian^26^ as

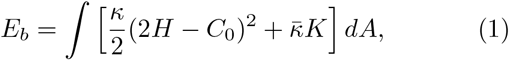

where *H, K*, and *C*_0_ are mean, Gaussian, and spontaneous curvatures, and *κ* and 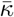 are the bending and Gaussian curvature moduli, respectively, and the integration is performed over the membrane area *A*. The higher-order terms in curvature are only relevant when the resolution of the description becomes comparable to the membrane thickness. It has been shown that Eq. 1 is suitable for length scales larger than 10 nm^66^.

To evaluate Eq. 1 numerically, surface discretization is needed. A popular and robust method is DTS. In this approach, the membrane surface is represented as a triangular mesh *M* = (*V, F, E*), where *V* = {**v**_1_, …, **v**_*m*_ } with **v**_*i*_ ∈ ℝ^3^ represents the vertex set, *F* = {*f*_1_, …, *f*_*k*_} denotes the face set, and *E* = {*e*_1_, …, *e*_*n*_} comprises the edge set. Each face *f*_*p*_ = (*i, j, k*) consists of a tuple of three vertices that form a closed triangle, while each edge *e*_*q*_ = (*j, k*) is a tuple of two vertex indices corresponding to connected vertices in the mesh. In the context of membrane modeling, each vertex represents a membrane patch containing hundreds of lipids. Mesh properties such as mean and Gaussian curvature (Eq. 1) can be obtained using different approaches^67–69^. Here, DTS simulations were performed using FreeDTS (v2.0, 24), which internally uses the shape operator approach^70^.

During simulation, both vertex positions and mesh connectivity evolve through Monte Carlo moves, standard vertex update, and Alexander move (edge flip), allowing the system to sample different membrane configurations while maintaining a physically realistic surface. Moreover, inhomogeneity can be induced into the model through the concept of inclusion, which can be used to model membrane domains and membrane proteins^24^. The acceptance probability of a proposed membrane configuration is given by the Metropolis-Hastings criterion^71,72^

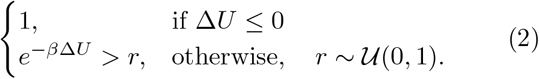

Here, Δ*U* is the total energy change between the proposed and current state, *β* = 1*/k*_*b*_*T* is the inverse temperature, and *r* is a number drawn from a uniform distribution.

While DTS simulations effectively model membrane physics based on theoretical principles, they traditionally lack the ability to incorporate experimental observational data. This disconnect between theory and experiment means that standard DTS simulations may not accurately capture the specific membrane configurations observed in real biological systems due to important hidden or unknown shape remodeling drivers. Our HMFF approach addresses this fundamental limitation by directly integrating experimental density data with physical simulation, creating a bridge between observed structures and the underlying biophysical principles.

### B. Helfrich Monte Carlo Flexible Fitting

HMFF extends the mesoscale DTS framework by incorporating volumetric experimental data, such as 3D EM, directly into simulations, creating a bidirectional feedback between physical modeling and experimental evidence. In practice, HMFF is implemented as an additional energy term that guides vertices towards experimentally observed membrane configurations, specifically regions of high electron density in 3D EM data. Simultaneously, the physical membrane properties included during DTS simulation (see methods IV A) act as regularizers on the ensemble of possible membrane configurations.

The guidance energy term used in HMFF is adapted from the approach introduced for Molecular Dynamics Flexible Fitting (MDFF) of atomic structures to electron density maps^27–30^

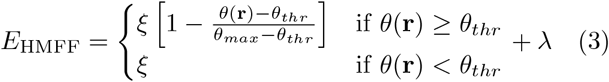

where *θ*(**r**) is density at position **r**, *ξ* controls interaction strength, *θ*_*max*_ the maximum value of the densities, and *θ*_*thr*_ the threshold value to exclude solvent contribution. In HMFF, *θ*_*thr*_ defaults to 0 and *θ*_*max*_ defaults to the 0.999 quantile of the potential map, which leaves *ξ* as the primary tunable parameter. Because neighboring vertices should occupy regions of similar density, we add a coherence regularization term per vertex

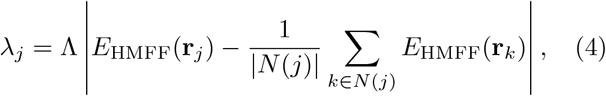

where *k* ∈ *N* (*j*) is a vertex in the 1-ring neighborhood of vertex *j* and Λ controls the regularization strength. In this work, we set Λ to 0 or 1 to match the scale of *E*_HMFF_, though other values are in principle possible. This term penalizes vertices whose HMFF energy departs from the local neighborhood mean, stabilizing the simulation against configurations where adjacent vertices end up in different density regions, particularly at high *ξ* where the data dominates. To extrapolate beyond boundaries of experimental volumes, we implement padding in the xy-plane using 0.4 × *θ*_*max*_ times the moving average within a three-slice window, while z-slices are padded using moving averages of terminal slices.

We implemented HMFF within FreeDTS as an additional energy term using the abstract energy class framework. Our implementation also includes a tri-linear interpolator for processing CCP4/MRC format potential maps at arbitrary spatial resolution. The HMFF term can be brought into action by using the parameter EnergyMethod FreeDTS1.0 HMFF. FreeDTS is available at https://github.com/weria-pezeshkian/FreeDTS, and HMFF exists in version 2.1 and above. We provide an overview of all energy terms available in FreeDTS in section X B.

### C. The Mosaic software

To address the challenges in practical HMFF implementation with experimental data, we developed Mosaic, a software platform for building models amenable to physical simulation on the scale of entire cells. Implemented as a Python-based application with a PyQt6/VTK^73,74^ graphical user interface (GUI), Mosaic combines existing tools with novel computational methods to enable the complete workflow from 3D EM data to physical simulations.

Mosaic uses MemBrain-seg^19^ for deep-learning-based membrane segmentation, the output of which is refined, clustered, and analyzed within the GUI. Mosaic converts segmentations into triangular meshes using a biologically motivated meshing approach, which are subsequently equilibrated to be suitable for simulation. Equilibrated meshes can be used in DTS simulation using FreeDTS^24^ with the HMFF potential. Mosaic also provides a dedicated interface for setting up DTS parameter screens and analyzing the resulting trajectories (e.g., Fig. 1c-g, 2d, 5c). HMFF meshes can be morphometrically characterized within Mosaic, used in equilibrium simulations, or form the basis for template matching of membrane proteins using PyTME^33^. Mosaic implements a dedicated projection procedure, which allows for analyzing membrane-protein characteristics, such as distribution and curvature preference. The combined HMFF mesh and protein picks can be mapped back onto coarse-grained representations using TS2CG^34,35^, enabling analyses from mesoscale dynamics to molecular interactions. All analyses presented in this work were performed through Mosaic, which is available at https://github.com/KosinskiLab/mosaic.

### D. Creating meshes for DTS simulation

Mosaic converts segmentations into meshes for DTS simulation in a multi-stage process: initial construction of a mesh from segmentation data, completion of the mesh to form a closed surface representing a biologically plausible membrane, and regularization to ensure mesh properties suitable for DTS simulation.

#### 1. Mesh construction

Mosaic implements complementary methods for creating meshes from segmentations, that is, finding a mesh *M* that approximates the surface formed by an unordered set of points *P* = {**p**_1_, …, **p**_*n*_} ∈ ℝ^3^. Each approach offers distinct advantages, with overall applicability depending on data characteristics. Mosaic provides optimized default parameters for each method derived from testing across diverse biological datasets.

Ball Pivoting^75^ provides robust mesh construction and only requires tuning a single parameter, the pivoting radius, which determines the maximum local curvature that can be captured. The pivoting radius is limited by the sampling density of the point cloud, i.e., no sub-voxel precision mesh construction is possible. An alternative is Poisson reconstruction^76^, which fits an implicit function to generate smooth surfaces. This method can handle variable feature sizes but requires reliable normal vector estimates and more careful parameter tuning for accurate mesh construction. *α*-shapes^77^ are suitable for segmentations with large topological gaps and can effectively bridge discontinuities while preserving the overall structure. Reducing the *α* parameter allows for capturing concave features such as membrane invaginations, though excessively small values can lead to disconnected components. Finally, for densely labeled volumetric data, e.g., the FIB-SEM-based segmentations of entire mitochondria as opposed to just their membranes, Marching Cubes^78^ efficiently constructs meshes at grid resolution, but requires parameter tuning to avoid erroneous merges or inaccurate representation in high-curvature regions. See section X A for a mathematical description of the individual algorithms and Fig. S9 for an illustration of the different use-cases of each.

#### 2. Mesh completion and deformation

The mesh creation methods outlined above are not guaranteed to result in a closed surface that is a realistic representation of biological membranes. Therefore, we devise targeted completion strategies, which are outlined below and illustrated in Fig. S10.

For meshes with identifiable hole boundaries, Liepa triangulation^79^ systematically closes holes by identifying boundary loops and applying iterative subdivision. The subdivision process minimizes the ratio between edge length and geodesic distance until hole closure

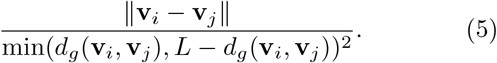

*d*_*g*_(**v**_*i*_, **v**_*j*_) represents the geodesic, ∥**v**_*i*_ − **v**_*j*_∥ the Euclidean distance along the boundary and *L* is the total boundary edge length. This process continues until each subdivision forms a triangle, effectively closing the hole while maintaining the overall shape. For meshes without a connected set of boundary vertices, we utilize *α*-shapes. After constructing the initial mesh, we compute distances between the original point cloud and mesh vertices. Regions, where distances exceed a threshold, by default 6× voxel size, are identified as missing areas requiring geometric optimization (Fig. S10a).

Both completion strategies introduce inferred vertices into the mesh. To reconcile inferred local and observed global geometries, we smooth the mesh using biologically motivated polyharmonic deformation (Fig. S10b)

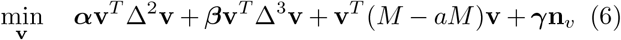

subject to Dirichlet boundary conditions **v** |_∂Ω_ = **v**_real_^80^, which fixes the positions of non-inferred vertices. Δ ∈ ℝ^*n*×*n*^ represents the discrete Laplacian operator, *M* ∈ ℝ^*n*×*n*^ is the area-mass matrix derived from Voronoi tessellation, and **n**_*v*_ is the normal of vertex **v**. The weights ***α, β, γ*** ∈ ℝ^*n*^ control smoothness, curvature, and pressure respectively where

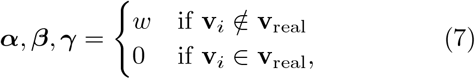

where *w* ∈ ℝ is a user-defined factor for each weight.

The biharmonic term ***α* v**^*T*^ Δ^2^**v** penalizes bending and promotes a smooth, *C*^1^-continuous surface, while the triharmonic term ***β* v**^*T*^ Δ^3^**v** imposes higher-order, *C*^2^ continuity for smooth curvature transitions between observed and completed regions. For further details on higher-order Laplacians in the context of triangular meshes, we refer the reader to Kobbelt *et al*. ^81^.

We construct Δ and *M* using intrinsic Delaunay triangulation with edge length mollification, which introduces a mollification factor for numerical stability

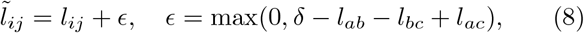

where 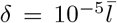 and 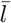 is the mean edge length. This ensures the triangle inequality is strictly satisfied in all mesh elements.

The Laplacian is then constructed using the cotangent formula as implemented in libigl^82^

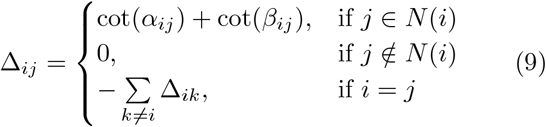

where *N* (*i*) are the vertices adjacent to vertex *i*, and *α*_*ij*_, *β*_*ij*_ are the angles opposite to edge *i, j*. The mass matrix is derived from Voronoi vertex areas, providing an appropriate inner product space for solving the polyharmonic system presented in Eq. 6.

#### 3. Mesh regularization

Not every mesh is suitable for DTS simulations, as the vertex update scheme and edge flips can result in a mesh that no longer represents a realistic membrane surface, for example, a surface intersecting itself. This demands that the edge length falls within a specific range 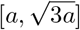 with *a* ∈ ℝ^+^ and that the dihedral angle between two adjacent triangles satisfies cos(*θ*) *>* − 1^24^, conditions that enforce self-avoidance throughout the simulation.

We devised a two-step equilibration process to create meshes suitable for DTS simulation. First, isotropic explicit remeshing^83^ is applied to adjust vertex positions and connectivity. The edges of the resulting mesh are subsequently regularized by minimizing the following energy term using Trimem (v0.2.2, 25)

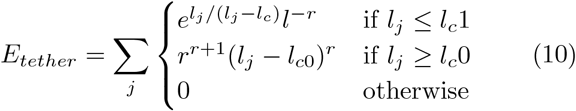

where *l*_*j*_ is the length of edge *j, l*_*c*_1 and *l*_*c*_0 being the lower-and upper-onset of penalization and *r* the slope of the potential well. Constraining the equilibration to preserve surface area, volume, and local curvature ensures that all edge lengths converge to the desired range while maintaining the overall shape of the mesh.

### E. Mesh projection

In biological systems, numerous entities of interest, from individual proteins to contact sites with membranes of other organelles, exist in relation to membrane meshes. To accurately characterize membrane-associated structures, mesh projection is required to relate spatial orientation to the continuous mesh representations. The projection process serves dual purposes: First, it enables rigorous structural analysis, e.g., quantification of protein curvature preference, geodesic distance distributions, and co-localization patterns. Second, projection ensures that *in silico* model systems, e.g., for equilibrium DTS or MD simulation, faithfully recapitulate the experimentally observed protein positions. Mosaic includes a set of computational methods to project objects onto membrane meshes and quantify their geometric context, which are described below and illustrated in Fig. S5d.

#### 1. Dual-mode projection

Mosaic implements a ray-casting-based dual-mode projection algorithm for mapping structures (e.g., protein positions, organelle segmentations, membrane meshes) onto triangulated mesh surfaces. The first mode computes the closest point on the mesh by minimizing Euclidean distance

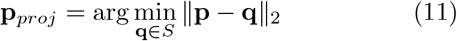

where *S* is the surface defined by the mesh, **q** ∈ ℝ^3^ a position on *S*, and **p** ∈ ℝ^3^ is the position to be projected.

The second mode accounts for structures with defined position and orientation, such as proteins relative to a membrane surface. On the level of individual points, they can be defined as a tuple (**p** ∈ ℝ^3^, *R* ∈ SO(3)). Given the principal axis of the protein, which Mosaic internally sets as *R***e**_*z*_, we modify the projection procedure to cast rays from the position along the principal axis

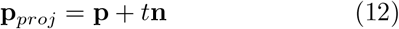

where *t* ∈ ℝ_≥0_ is the distance to the first intersection with the mesh surface along **n** (Fig. S5d). Particularly in regions of high curvature or for proteins that are bent from the mesh surface normal, the second mode is more accurate.

#### 2. Mesh extension

To obtain precise geometric properties at projected positions, we extend the mesh by incorporating these positions as new vertices. Given a set of projected points that intersect with a triangle *f*_*i*_ formed by vertices (**v**_*a*_, **v**_*b*_, **v**_*c*_) ∈ ℝ^3^, we map them into the local 2D reference frame of *f*_*i*_ as

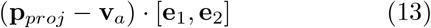

where **e**_1_ and **e**_2_ form an orthonormal basis in the triangle plane. A valid surface triangulation of this 2D space that includes all projection points is obtained using Delaunay triangulation implemented in scipy.

The extended mesh preserves the original surface geometry while including exact representations of projected positions, enabling more accurate computation of local properties compared to interpolation approaches using barycentric coordinates.

#### 3. Mesh property computation

Mesh properties such as curvatures and geodesic distances can be computed on the extended mesh. Curvature computation uses the local vertex n-ring neighborhood with radius *r* to estimate principal curvatures *k*_1_ and *k*_2_ at each vertex^82^. Based on principal curvatures, the Gaussian curvature *K* and the mean curvature *H* can be computed as

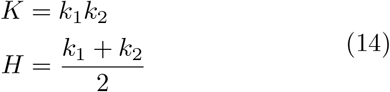

Geodesic distances between vertices on the mesh surface are computed using an exact algorithm^84^, which provides accurate surface distances even in regions of high curvature where Euclidean measurements would be inaccurate.

### F. Mycoplasma pneumoniae data acquisition and processing

*M. pneumoniae* cryo-ET data was collected and processed as previously described^38^. In brief, the bacterium was cultured on gold EM grids with a holey carbon film (Quantifoil) before being vitrified by plunge freezing. The sample was imaged on a Titan Krios G3 microscope (Thermo Fischer) equipped with a K2 direct electron detector (Gatan), dose symmetrically between -60 and 60 degrees with 3-degree increments at a pixel size of 1.7 Å. Tilt series processing was performed using Warp, and the initial tilt series alignment was determined using IMOD^85,86^. A ribosome reference was created from manually picked ribosomes in RELION at a voxel size of 6.082 Å. It was subsequently low-pass filtered to 30 A and used directly as a template in PyTOM^87^. The 400 highest-scoring cross-correlation peaks were extracted and filtered by visual inspection. Ribosomes were subtomogram averaged using RELION, and used as fiducials for improving tilt series alignment in M^88,89^. The updated alignments were used to reconstruct the final tomograms in Warp.

### G. Membrane segmentation

Membranes of *M. pneumoniae* and IAV VLPs (EMD:11075, 51) tomograms were segmented using MemBrain-seg (v0.05, 19), with Mem-Brain seg v10 alpha weights, which were downloaded from https://github.com/teamtomo/membrain-seg. The input tomograms were resampled to a voxel size of 12 Å using the utilities provided with MemBrain-seg. Segmentation was performed using 8-fold test time augmentation and a 160-voxel sliding window on an NVIDIA RTX 4070 Super GPU.

### H. Initial mesh generation

The initial mesh for the planar membrane simulation in a potential field was generated using the GEN script provided within FreeDTS (v2.0, 24). The corresponding potential map was generated as an isotropic Gaussian with *σ* = 4, centered ten voxels above the planar membrane mesh using PyTME (v0.3.0, 33).

Initial meshes for *M. pneumoniae* were generated using Mosaic. Membrane segmentations were cleaned by removing erroneous segments and thinning the segmentations to the outer cloud. The cleaned segmentations were triangulated using *α*-shapes with an elastic weight of 1.0 and a curvature weight of 10.0. We achieved a more realistic estimate of cell volume by performing 100 Å equidistant sampling from the created mesh, manually removing undesirable samples, and repeating the *α*-shape triangulation with a pressure of 0.1. The resulting triangulation was remeshed to an average edge length of 170 Å and equilibrated using default parameters (bending energy coefficient 300.0, area conservation coefficient 10^6^, volume conservation coefficient 10^6^, edge tension coefficient 10^5^, surface repulsion coefficient 10^3^), with volume and area fraction targets of 1.1, ensuring that overall shape is maintained while creating a mesh suitable for dynamic triangulated surface simulations. Initial meshes of IAV were created analogously to *M. pneumoniae* but remeshed to an average edge length of 110 Å and 70 Å.

Initial meshes of cellular organelles were generated using publicly available segmentations^22^. EM data, organelle segmentations, and ribosome positions for jrc-hela2 were downloaded from https://open.quiltdata.com/b/janelia-cosem-datasets/tree/jrc_hela-2/ using custom Python scripts. Initial meshes were generated from the unbinned s0 data layer, providing a resolution of (4 nm, 4 nm, 5.24 nm) along *x, y, z*. One-time binned data was used specifically for the ER, as it yielded visually higher quality meshes. Volumes were converted into triangular meshes by marching cubes. Briefly, volumes were split into non-overlapping subvolumes with a box size of 448 and meshed separately using zmesh (v1.8.0, https://github.com/seung-lab/zmesh). The initial meshes were obtained using a reduction factor on a triangle count of 100 and a maximum simplification error of 40 nm. Following simplification, meshes were merged and simplified again using pyfqmr (0.3.0, https://github.com/Kramer84/pyfqmr-Fast-Quadric-Mesh-Reduction), primarily to reduce elevated triangle count at volume boundaries. The second simplification was performed using an aggressiveness of 5.5 and a decimation factor of 2.0. These settings are the default in Mosaic, which uses an implementation inspired by Igneous^90^.

Meshes of cellular organelles were imported into Mosaic and assessed for quality. Meshes were repaired by equidistant sampling from the surface, closing gaps using face-projection and equidistant sampling via *α*-shapes, and manually removing erroneous segmentations. Subsequently, cleaned point clouds were meshed using Poisson reconstruction, using default Mosaic parameters but varying the depth between 9 and 14 depending on the complexity of the point cloud. Mitochondria meshes were prepared for DTS simulation analogous to previous examples, but remeshed to 20 nm.

### I. HMFF simulation

Tomograms of *M. pneumoniae* and IAV VLPs were obtained at a voxel-size of 13.6 Å and 6.8 Å, respectively. Through Mosaic, they were bandpass filtered to 900 Å and 50 Å for IAV and 900 Å and 140 Å for *M. pneumoniae*. The latter were further normalized by dividing each z-slice by its maximum density value. For jrc-hela2, we used a volume with voxel size of (16 nm, 16 nm, 20.96 nm) and no further preprocessing. HMFF simulations were run on an AMD Ryzen 5 7600 using 32GB of DDR5 RAM. An overview of key parameters used for HMFF simulation is presented in Tab. 1.

**TABLE 1.**
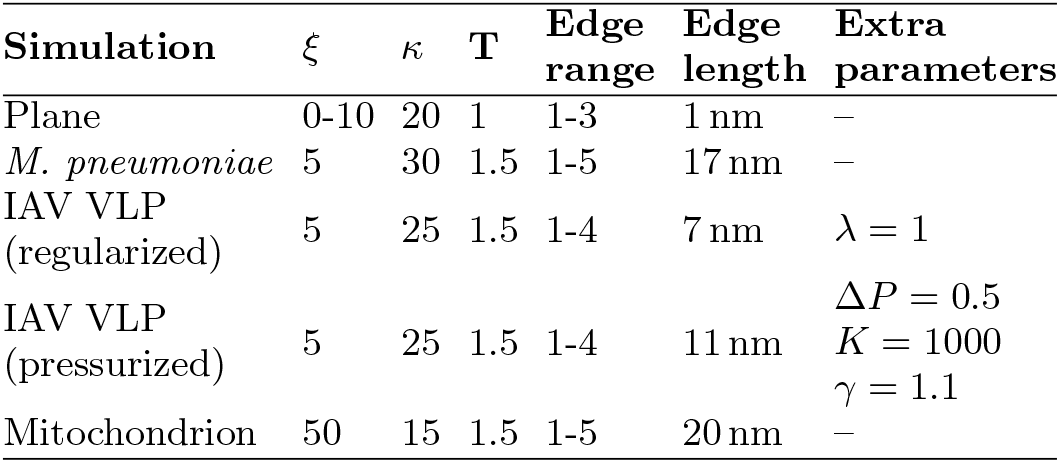
Overview of key HMFF simulation parameters. Edge range is the permitted edge-length interval in DTS units; edge length is the physical average. Extra parameters lists run-specific terms: volume coupling via the polynomial potential 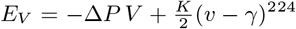, and coherence regularization with weight *λ*. The elevated *κ* for *M. pneumoniae* reflects its high cholesterol content^**?**^ .

#### 1. Undulation spectrum

For each frame of the planar-membrane trajectory, vertex positions were projected onto the *xy* plane and the height field *u*(*x, y*) obtained by linear interpolation of the vertex *z* coordinates onto a regular 64 × 64 grid spanning the periodic box. After subtracting the mean height, we computed the power spectrum |*û*(**q**)| ^2^*/A* from the discrete Fourier transform *û*(**q**), with *A* = *L*_*x*_*L*_*y*_ the projected area. The radially averaged spectrum ⟨|*u*(*q*)|⟩ ^2^ was obtained by binning into logarithmically spaced shells of *q* = |**q**|, excluding the zero mode and wavevectors above half the maximum sampled magnitude to avoid aliasing, and averaging over the trajectory (4.5 million steps after discarding the first 500 thousand steps). We compared the result to the Helfrich prediction for a tension-less membrane

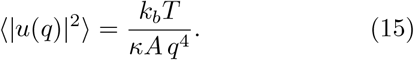

#### 2. Synthetic validation

Three synthetic reference systems with known physical drivers were simulated using FreeDTS and used to validate HMFF.

Sphere-to-dumbbell evolved a spherical membrane into a dumbbell shape with *κ* = 60, edge length 5, volume coupling target 0.7 (*k* = 10^4^), curvature coupling *C*_0_ = 0.3 (*k* = 150), and area coupling target 0.34 (*k* = 1000)^34^. Sphere-to-discoid used *κ* = 60, edge length 5, and volume coupling target 0.2 (*k* = 10^4^)^24^. Budding (BUD) was a planar membrane with *κ* = 20, edge length 10, simulated under isotropic frame tension and containing protein inclusions at density 0.15. Inclusions imposed spontaneous curvature (*C*_0_ = − 0.2, *κ*_inc_ = 20, 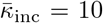) and an attractive inclusion-inclusion interaction (*n* = 1, Θ_0_ = 2, *C* = 4, *γ*_0_ = − 4)^24^.

Reference frames were converted to simulated densities by voxelization with trimesh at a voxel-size of 5 Å, Gaussian low-pass filtering to 40 Å, peak normalization to 1.0, and adding Gaussian noise with zero mean and standard deviation obtained as the ratio of signal in the membrane region over the requested SNR.

HMFF was run against these densities, in each case omitting a physical driver that had produced the reference shape, to assess whether the data alone could recover it. For the dumbbell and discoid, we disabled volume coupling and started HMFF from a sphere. Simulations were run over 5 million steps in three replicates with different random seeds. For the bud, we instead omitted the protein inclusions that drove budding and started from the budded reference at step 400 thousand. The HMFF simulation was run for 1 million steps.

#### 3. Experimental validation

*M. pneumoniae* cryo-ET data was processed as outlined above. Manually annotated membrane segmentations^20^ were imported into Mosaic and processed through a uniform pipeline. Segmentations were downsampled by center-of-mass clustering with radius 120 Å. Meshes were generated by shrink-wrap fitting^91^ with target edge length 100 Å, smoothness weight 1.0, 15 nearest-neighbor connectivity, gap bridging enabled, and 100 fitting iterations. Resulting meshes were equilibrated to an average edge length of 140 Å with parameters otherwise identical to those above.

HMFF was run with low-pass = 140 Å, high-pass = 900 Å, *κ* = 30, *ξ* = 7.5, and *λ* = 1. Each simulation ran for 3 million steps.

We validated HMFF through global and local data ablation, replacing voxels with values drawn at random from the same tomogram to preserve its noise characteristics. We report the average RMSE and its standard deviation between the manual segmentation and the HMFF mesh over the final 50 thousand simulation steps. In the global ablation, a fraction *p* of voxels, selected without replacement, was replaced. We swept *p* from 0 to 1 in steps of 0.05 and fit the RMSE to *a* exp(*bp*) + *c* by least squares. In the local ablation, rectangular masks of fixed *z*-height (80 voxels) and doubling width (10, 20, 40, 80, 160 voxels) were placed at manually selected concave, convex, and flat membrane segments.

#### 4. Error metric

Given a set of reference points *P* = {**p**_1_, …, **p**_*N*_}, corresponding to a segmentation or the vertices of a reference mesh, we computed distances to a given mesh *S* using ray-casting (projection mode 1 per section IV E).

We quantify the overall deviation using the Root Mean Square Error (RMSE) as

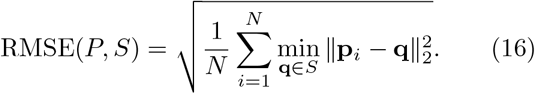

### J. Constrained template matching

Seed points for constrained template matching using PyTME^33,44^ were created from HMFF meshes using Poisson disk sampling^76^ with five-fold oversampling in Mosaic. *M. pneumoniae* seed points had an average Euclidean distance of 80 Å and were offset by the surface normal of the mesh by 80 Å. IAV seed points had an average Euclidean distance of 40 Å and were offset by 100 Å. These parameters reflect the expected glycoprotein spacing and positioning of membrane surface protein centers.

PDB:8pbz (Nap complex from *Mycoplasma genitalium*) was used as a template to pick Nap complexes in *M. pneumoniae*^43^. The corresponding structure was aligned to the z-axis, converted into a density map, low-pass filtered to 13.604 Å, and resampled using cubic spline interpolation. A spherical mask centered on the extracellular head group with a radius of 81.6 Å and smoothing sigma decay of 1.0 was used. For IAV, structural models of HA and NA from AlphaFold 2 multimer were used as templates^92,93^. HA models were based on the sequence of A/Hong-Kong/1/1968 H3N2 (UniProt: P11134), NA models on the sequence of A/California/04/2009 H1N1 (UniProt: C3W5S3). AlphaFold 2 multimer was run with default parameters except for increasing refinement cycles to 6. Structural models were converted to density maps analogously to *M. pneumoniae*. A cylindrical mask with a height of 251.6 Å, radius of 68.0 Å, and smoothing sigma decay of 2.0 was used for both templates. All templates and masks were created using PyTME^33^.

Template matching was performed using the FLC score, continuous wedge masks reflecting on the respective tilt range, and an angular sampling rate of 7 degrees. *M. pneumoniae* and IAV tomograms used for template matching had a voxel size of 6.80 Å. Matches were accepted when deviating no more than 15^°^ from the nearest seed point normal and falling within an ellipse of radii (10,10,20) voxels centered around that seed point and oriented along its normal. These cutoffs were changed for IAV to 15^°^ and radii of (6,6,10) and (6,6,12) for HA and NA, respectively. In *M. pneumoniae*, template matches were identified using PeakCallerMaximumFilter and a minimum peak distance of 20 voxels, a score cutoff of 0.09, and a distance to the HMFF mesh between 70.0 Å and 120.0 Å. For IAV, a minimum peak distance of 10 voxels and a score cutoff of 0.135 were used for HA, and a 7 voxel distance and 0.12 score cutoff were used for NA. Peaks were manually refined and filtered to be within 150.0 Å of the HMFF mesh for HA, and additionally no closer than 100.0 Å for NA. NA picks were filtered to include the top 97% quantile of scores, and HA picks falling within 7 voxels of remaining NA picks were removed to avoid clashes.

### K. Unconstrained template matching

*M. pneumoniae* ribosomes were picked using EMD:17132^39^ as template and a spherical mask with radius 142.8 Å. The initial template was low-pass filtered to 27.2 Å and resampled using cubic spline interpolation. Template matching was performed in PyTME as outlined above, but on a tomogram with voxel size 13.60 Å. Peaks were identified using PeakCallerScipy, a minimum peak distance of 15 voxels, and a score cutoff of 0.21. Picks were subsequently manually refined.

### L. Comparing membrane representations of the IAV VLP

We assessed the accuracy of the HMFF mesh against the cleaned membrane segmentation and a cylinder fit to the segmentation as an idealized representation of the complete VLP, using the HA picks from constrained template matching (IV J). We used the HMFF mesh refined without volume coupling, as the volume-coupled mesh from the main analysis informed constrained template matching. We used only HA picks, as the tip-localized NA picks would bias the comparison toward the cylinder’s flat caps.

All processing was performed in Mosaic. The membrane segmentation is a thick, voxel-resolution point cloud against which distances and angles are ill-defined. Therefore, we extracted its medial skeleton (skeletonization, method core and sigma 1.0, adapted from 94), denoised it by statistical outlier removal (20 neighbors, standard-deviation ratio 2.0), and downsampled it by center-of-mass binning (radius 60 Å) before meshing by Poisson reconstruction. The cylinder was fit to the same medial-skeleton point cloud by minimizing the squared residual and converted into a mesh using an *α*-shape (*α* = 1, equivalent to the convex hull).

We projected HA picks onto each mesh using closest-point projection (IV E, mode 1) rather than ray-cast projection (mode 2), because rays along the HA principal axis did not reliably intersect the incomplete segmentation. For each projection, we computed the Euclidean distance to the projected point and the angle between the pick’s principal axis and the surface normal at that point. We computed an expected distance between a pick and the membrane of approximately 9 nm, the offset between the center of mass and the centroid of the trans-membrane region (residues 528-552) in the HA structural model. We referenced the center of mass rather than the full structure to match the convention of our picks.

Since HA picks were identified using seed points derived from the HMFF mesh and manually refined (section IV J), we considered how this could bias the comparison. Template matching constraints permitted positional deviations up to 134 Å and angular deviations up to 21^°^ (following from the cone half-angle *θ* via *ϕ* ≤ arccos (cos^2^ *θ*), 44) in theory, relaxed by voxelization to 140 Å and 30^°^ in practice, relative to the mesh used for picking. Because the observed distance and angle distributions fell within these limits rather than accumulating at the cutoffs (Fig. S6b-c), we concluded that the acceptance window did not substantially truncate them.

### M. Backmapping DTS onto coarse-grained Martini models

TS2CG (v1.2.2, 34,35) was used to backmap DTS onto coarse-grained models using Mosaic. Briefly, HMFF meshes of *M. pneumoniae* were remeshed to 120 Å and IAV meshes to 20 Å. Picked proteins were projected onto the mesh (mode 2 per section IV E) and mapped to the closest vertex.

Atomic structures were coarse-grained using the martinize2 (v0.13.0, 95) tool with default parameters and using an elastic network with standard cutoffs to maintain the tertiary structure of NA, HA, and the Nap complex. TS2CG PCG was run assuming a bond length of 0.1 and a CG lipid library downloaded from https://github.com/weria-pezeshkian/TS2CG-v2.0^96^. For simplicity, a pure POPC bilayer was generated using a thickness of 3.8 nm and an area per lipid of 0.64 and 0.5 for *M. pneumoniae* and IAV, respectively. Proteins were inserted into the bilayer with a normal offset of 6.5 nm for the Nap complex, 7.5 nm for HA, and 11.5 nm for NA. Since TS2CG does not allow for including per-protein orientations, HA and NA particles identified by template matching clashed. Inter-protein clashes were identified, and those with a distance of *<* 0.3 nm were removed, followed by membrane rebuilding to avoid holes in the regions with removed proteins. Subsequent energy minimization and equilibration were performed using the GROMACS simulation package^46^. The system was equilibrated without explicit solvent following the TS2CG protocol^34^. In general, the system could be solvated using conventional tools, however. The structures were first minimized using the steepest descent algorithm with soft core potentials (init-lambda = 0.001, then 0.0005) to alleviate remaining steric clashes, followed by a 1000-step minimization using regular potential with reaction-field electrostatics. The minimized structure was then equilibrated with protein and lipid headgroup position restraints using a stochastic dynamics integrator with a 1 fs timestep and Berendsen thermostat set to 310 K. Pressure coupling remained deactivated throughout the equilibration to account for the missing solvent, while maintaining the Verlet cutoff scheme with periodic boundary conditions in all dimensions. For*M. pneumoniae*, soft core potentials were disabled, and the equilibration step was omitted. Mdp files with a complete list of settings can be found in the supporting files.

### N. Mitochondria classification

Three complementary feature sets capturing topological and geometrical properties were derived to differentiate latent mitochondria classes.

Topological information was captured through the spectral properties of the Laplace operator by solving the generalized eigenvalue problem Δ*ϕ*_*i*_ = *λ*_*i*_*Mϕ*_*i*_ for the first 15 eigenvalues with the smallest magnitude. These eigen-values encode the intrinsic topology of the mesh, which is independent of the spatial embedding. The sparse system was solved using scipy’s sparse eigenvalue solver (v1.14.1, 97) with shift-inversion *σ* = 10^−8^.

Geometric information was captured by 3D Zernike moments as rotation-invariant shape descriptors. The position of each mitochondrial vertex **v**_*i*_ ∈ *V* was normalized as (**v**_*i*_ − ***µ***)*/r*_max_, where 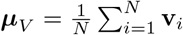 and *r*_max_ = max_*i*_ ∥**v**_*i*_ − ***µ***_*V*_∥_2_, ensuring unit sphere containment. 3D Zernike moments were computed from normalized vertices as

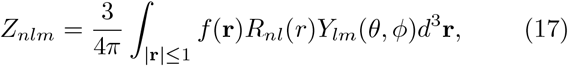

where *R*_*nl*_(*r*) are radial polynomials of order *n* with *n* − *l* being even and non-negative, *Y*_*lm*_(*θ, ϕ*) are spherical harmonics of degree *l* and order *m* with |*m*| ≤ *l, f* (**r**) is the point cloud density function defined within the unit sphere, and |**r**| is the position vector with magnitude *r* = |**r**|. In spherical coordinates, the volume element *dV* = *d*^3^**r** is expressed as *r*^2^ sin *θdrdθdϕ* with *θ* ∈ [0, *π*] and *ϕ* ∈ [0, 2*π*]. Zernike moments were computed using https://github.com/kheyer/3D-Zernike-Descriptors. The rotation-invariant descriptors 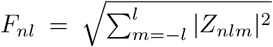, were calculated with *l* ranging from *l*_0_ = *n* mod 2 to *n* in steps of 2, yielding (*n*+1)(*n*+2)*/*2 descriptor coefficients. The descriptors were computed up to order *n* = 8.

Topological and geometric descriptors were supplemented with scale-dependent features: surface area, volume, skeleton branch count, and curvature statistics. Mosaic calculated surface area using Open3D (0.19.0, 42), volume using tetrahedron decomposition, and mean and Gaussian curvature using a radius of ten. Topological complexity was quantified through segment counts using skeletor (v1.3.0, https://github.com/navis-org/skeletor). A wavefront-based skeletonization algorithm was applied with 5 waves, and segments longer than 5 nodes were counted.

All features were standardized to a zero mean and unit variance. Principal component analysis was applied to the standardized features, followed by t-SNE dimensionality reduction using Rtsne (v0.17, 98) with parameters perplexity 10, early exaggeration 24, max iterations 15,000, and random initialization. A nearest-neighbor graph was constructed from the principal-component embedding, and Louvain community detection was performed on this graph using igraph (v2.0.3, 99) with a resolution parameter of 1.0. To assess whether the resulting clusters depended on this particular t-SNE configuration, we repeated the embedding following the protocol of Kobak and Berens^100^, implemented in OpenTSNE and accessed through snifter (v1.18.1, 101). As this protocol was developed for single-cell RNA sequencing data, whose scale differs markedly from ours, we adopted its settings but changed the perplexity to 10, since the prescribed value scaled to our dataset is degenerately small. Cluster assignments were preserved under this alternative protocol, the only meaningful difference being the merging of clusters 2 and 3 (Fig. S8d).

### O. Contact point analysis

Contact points were determined using distance-based detection. Geometric proximity was computed using Mosaic, which internally relies on the Open3D^42^ ray-casting scene implementation. For each mitochondrion mesh and target organelle segmentation, spatial relationships were analyzed. A contact was defined when any part of the organelle segmentation was located within 14 nm of the mitochondrial surface, similar to the threshold used in previous analyses of this dataset^22^.

The contact area was calculated by summing the triangular face areas from the mitochondrial mesh that fell within the contact threshold. The percentage of mitochondria within each morphological cluster forming contacts with other organelles was determined by counting mitochondria with non-zero contact area, then dividing by the total number of mitochondria in each respective cluster.

## V. DATA ANALYSIS

Three-dimensional renderings of meshes, tomograms, and molecular systems, as well as the metrics reported throughout (e.g., curvatures, geodesic distances, contact areas, and RMSE), were produced in Mosaic using the implementations described above. Backmapped coarse-grained models were visualized using VMD (v1.9.3)^102^. Data analysis and visualization were performed in R (v4.5.0)^103^ using ggplot2^104^ (v3.5.2), data.table^105^ (v1.17.0), ggpubr^106^ (v0.6.0), ComplexHeatmap^107^ (v2.24.0) and ggdist^108^ (v3.3.3).

## VI. DATA AVAILABILITY

Mosaic sessions, intermediate results, source code, and cryo-ET data of *M. pneumoniae* will be deposited in Zenodo and are available from https://oc.embl.de/index.php/s/iGPdEJGfoXVhSno for peer review.

## VII. CODE AVAILABILITY

HMFF is available in FreeDTS from https://github.com/weria-pezeshkian/FreeDTS. Mosaic is available from https://github.com/KosinskiLab/mosaic.

## VIII. ACKNOWLEDGEMENTS

We thank the EMBL IT and HPC resources for providing essential computational infrastructure. VM and JK acknowledge funding from the CSSB flagship project Plasmofraction. MS acknowledges support from a research fellowship from the EMBL Interdisciplinary Post-doc (EIPOD) Programme under Marie Curie Cofund Actions MSCA-COFUND-FP (grant agreement number: 847543). RKJ was supported by the Independent Research Fund Denmark (grant number 0164-00010A). JM acknowledges the support from the EMBL, an EMBL Infection Biology Transversal Theme Synergy grant, and a Chan Zuckerberg Initiative grant for Visual Proteomics (grant No. 2021-234620). VM, JK, and JM were supported by the ERC (TransFORM, 101119142). WP acknowledges support from the Novo Nordisk Foundation (grant No. NNF18SA0035142 and NNF22OC0079182) and Independent Research Fund Denmark (grant No. 10.46540/2064-00032B).

## IX. COMPETING INTERESTS

The authors declare no competing interests.

## X. EXTENDED DATA

### A. Meshing Algorithms

Several established algorithms exist for generating triangular meshes from point clouds, each with distinct mathematical foundations and operational characteristics. The paragraphs below outline the approaches implemented in Mosaic and used throughout this work.

The Marching Cubes algorithm discretizes ℝ^3^ into a regular grid of cubes and constructs the surface through local triangulation. For each cube intersected by the implicit surface *f* (*x, y, z*) = 0, the algorithm determines the surface-edge intersections based on the sign of *f* at the cube vertices. The local triangle configuration is selected from a predefined set of 2^8^ possible cases, ensuring consistent topology between adjacent cubes. This produces a triangulation that approximates the level set of the implicit function^78^.

Ball Pivoting Algorithm (BPA) operates through a geometric rolling process. Starting from a seed triangle, a ball of radius *ρ* pivots around an active edge until it touches another point, forming a new triangle. Mathematically, this process identifies points **p** satisfying

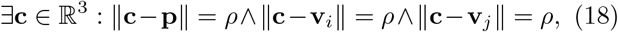

where **v**_*i*_, **v**_*j*_ are the vertices of the active edge. This generates a subset of the *ρ*-regular triangulation of the point set^75^.

*α*-shapes provide a formal mathematical framework for shape reconstruction through a filtration of the Delaunay triangulation. For a given *α* ∈ ℝ, the alpha complex consists of all Delaunay simplices whose dual Voronoi cells intersect an *α*-ball centered at each vertex. The resulting shape captures topological features at the scale determined by *α*, with the family of *α*-shapes forming a hierarchical representation of the point set^77^.

Poisson Reconstruction estimates the indicator function *χ* of the shape by solving the Poisson equation

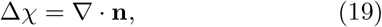

where **n** represents the oriented normal field sampled at the input points. The solution is efficiently computed using a multi-scale approach in an adaptive octree, with the final surface extracted as an appropriate level set of the resulting scalar function^40^.

### B. Energy terms of the FreeDTS framework

HMFF is implemented within FreeDTS^24^ and adds a data-derived term *E*_HMFF_ (see methods IV B), which guides membrane vertices toward regions of high density in the experimental data. FreeDTS expresses the energy of a triangulated mesh as a modular sum of contributions. Each term can be enabled independently, so that the Hamiltonian for a given system comprises only those terms relevant to it. Below, we summarize the categories of energy terms documented for FreeDTS and refer the reader to Pezeshkian and Ipsen ^24^ for explicit functional forms.

- *Local membrane mechanics*. The Helfrich bending energy, parametrized per vertex by the bending and Gaussian moduli and the spontaneous curvature; a line energy for vertices on open (non-closed) edges, comprising line tension and geodesic/normal curvature terms; and a local area-elasticity term penalizing deviations of per-vertex area (or per-edge length for edge vertices) from a target value.
- *Global geometric constraints*. Couplings that restrain extensive quantities of the whole mesh: enclosed volume (via a second-order polynomial potential or an osmotic-pressure term), total membrane area, and the area-integrated mean curvature, the latter capturing monolayer-area-difference (asymmetry) effects. For membranes that are periodic in one or more directions, a frame tension can additionally be imposed by coupling the simulation box dimensions to the mesh.
- *Inclusions and fields*. Membrane-embedded inclusions represent proteins or compositional domains and carry their own bending parameters and (optionally anisotropic) spontaneous curvature, together with inclusion-inclusion interaction potentials; in-plane vector fields and external fields can be applied to model orientational order and directed forces.
- *Environment and restraints*. Adhesion of vertices to a defined substrate, confinement within rigid boundaries (parallel walls or ellipsoidal shells/cores), and harmonic restraints between user-defined vertex groups, as well as bonded and nonbonded vertex potentials.

**TABLE S1.** Number of mitochondria per morphological cluster identified by Louvain community detection.

| Cluster | $N$ |
| --- | --- |
| 1 | 35 |
| 2 | 25 |
| 3 | 17 |
| 4 | 30 |
| 5 | 31 |
| 6 | 34 |
| 7 | 34 |
| 8 | 14 |
| 9 | 17 |

**Fig. S1.**
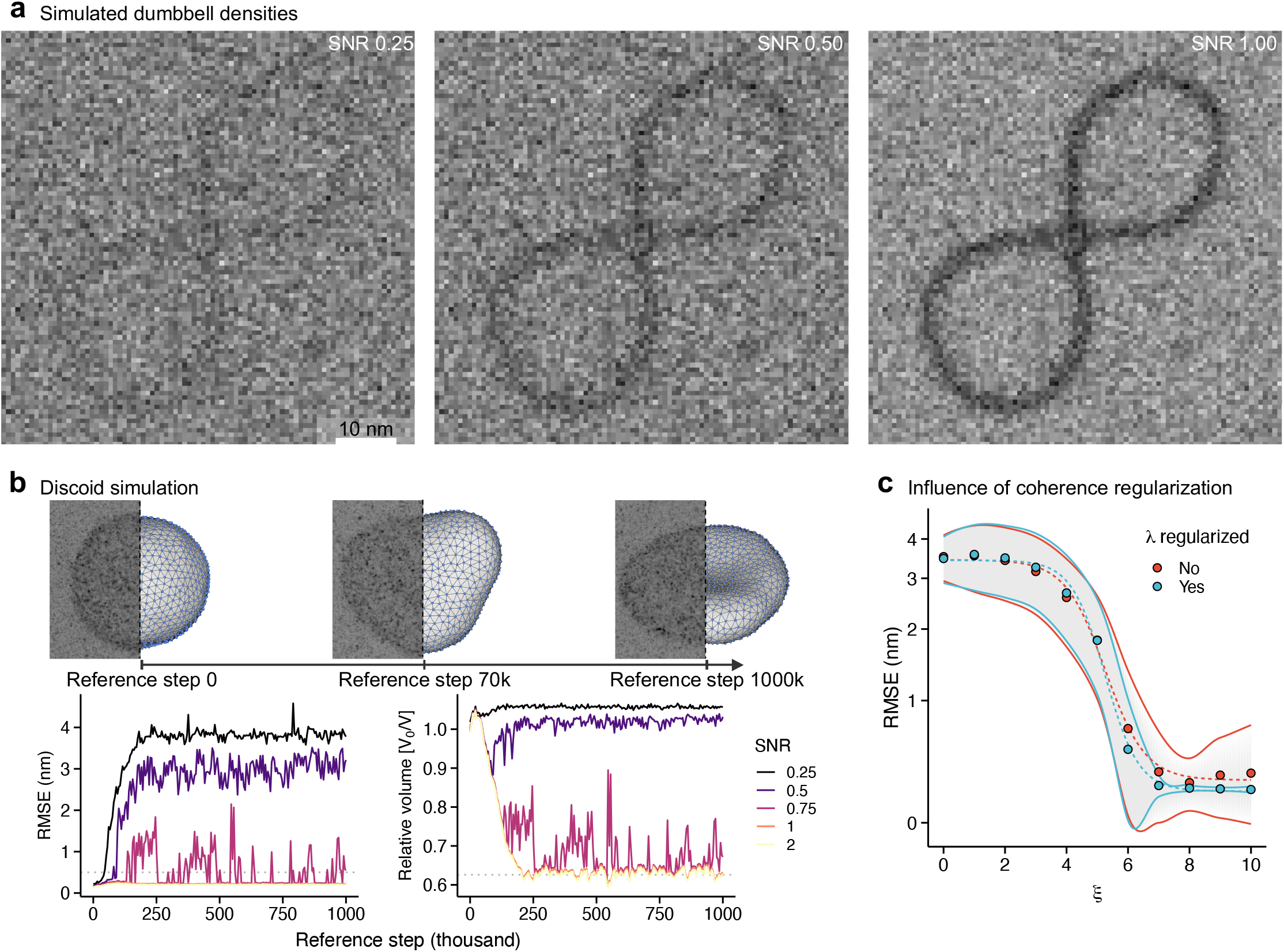
Validation of HMFF on synthetic data. **a**, Simulated densities of the dumbbell reference trajectory in Fig. 1d at the indicated SNR. **b**, Membrane simulation under volume coupling (top). Corresponding simulated density at SNR 0.5 shown as MIP. HMFF simulations started from a sphere and were run against simulated densities of each reference step at the indicated SNR with *ξ* = 4 and *λ* regularization. RMSE to the reference (left) and relative volume (right), averaged over 50k simulation steps and three replicates; dashed lines mark the voxel size of the simulated density and the realized volume ratio of the reference trajectory. **c**, Mean RMSE over all reference steps for the simulation in **b** versus coupling strength *ξ* at SNR 0.5, with and without *λ* regularization. Shaded bands indicate standard deviation.

**Fig. S2.**
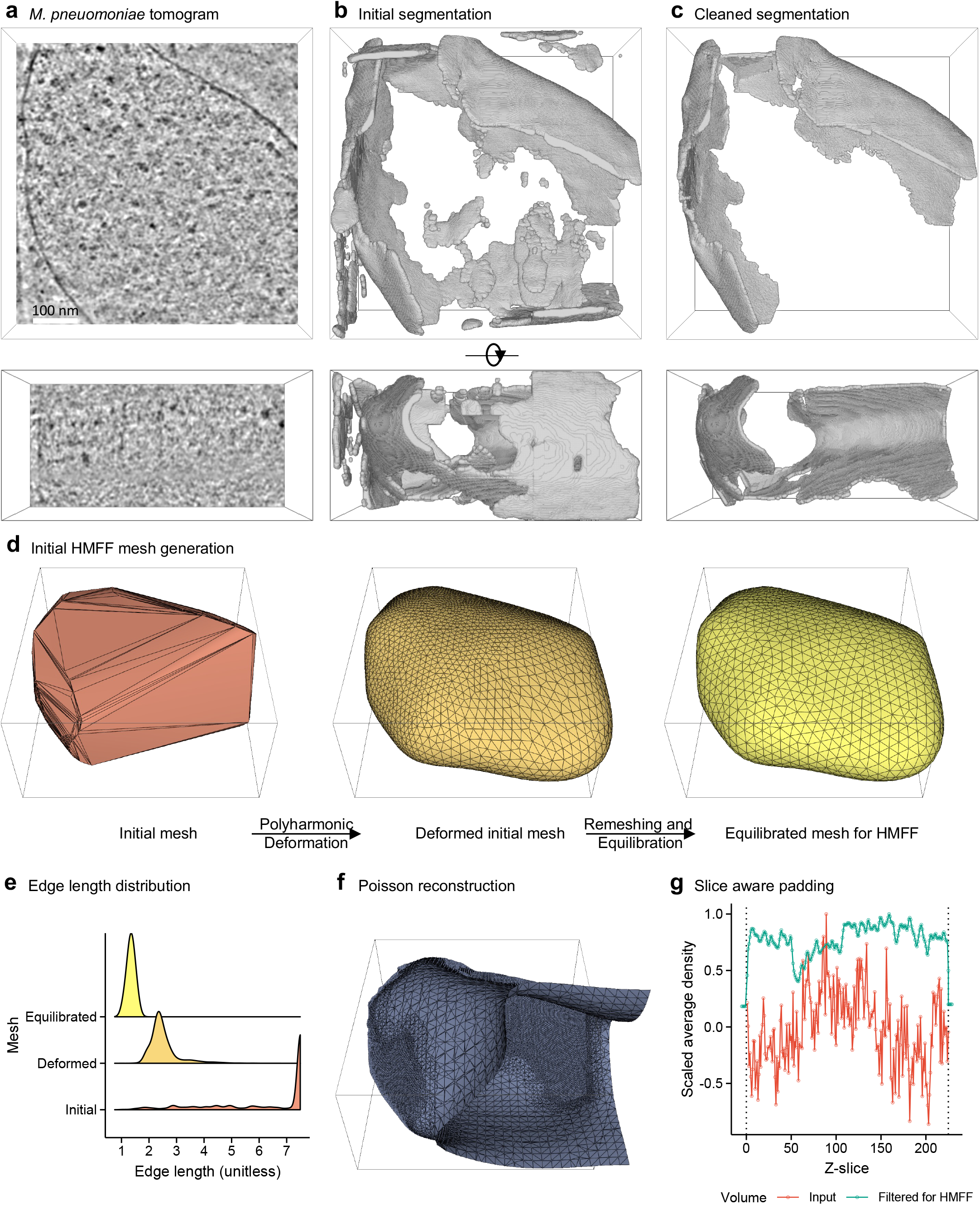
Setting up *M. pneumoniae* HMFF simulations. **a**, Band-pass filtered tomographic slice showing a *M. pneumoniae* cell. **b**, Initial membrane segmentation. **c**, Cleaned segmentation following processing in Mosaic. **d**, Creating a cell membrane mesh model suitable for HMFF simulation by polyharmonic deformation and subsequent equilibration of an *α*-shape fitted to refined membrane segmentation. **e**, Mesh edge lengths stratified by stage shown (**d**). **f**, Poisson reconstruction based on refined membrane segmentation. **g**, Padding values used by HMFF for the particular tomogram shown in (**a**), compared to the raw tomographic densities. Tomogram boundaries are indicated as vertical lines. Scaled average density corresponds to the per-slice average, divided by the maximum average value over all slices.

**Fig. S3.**
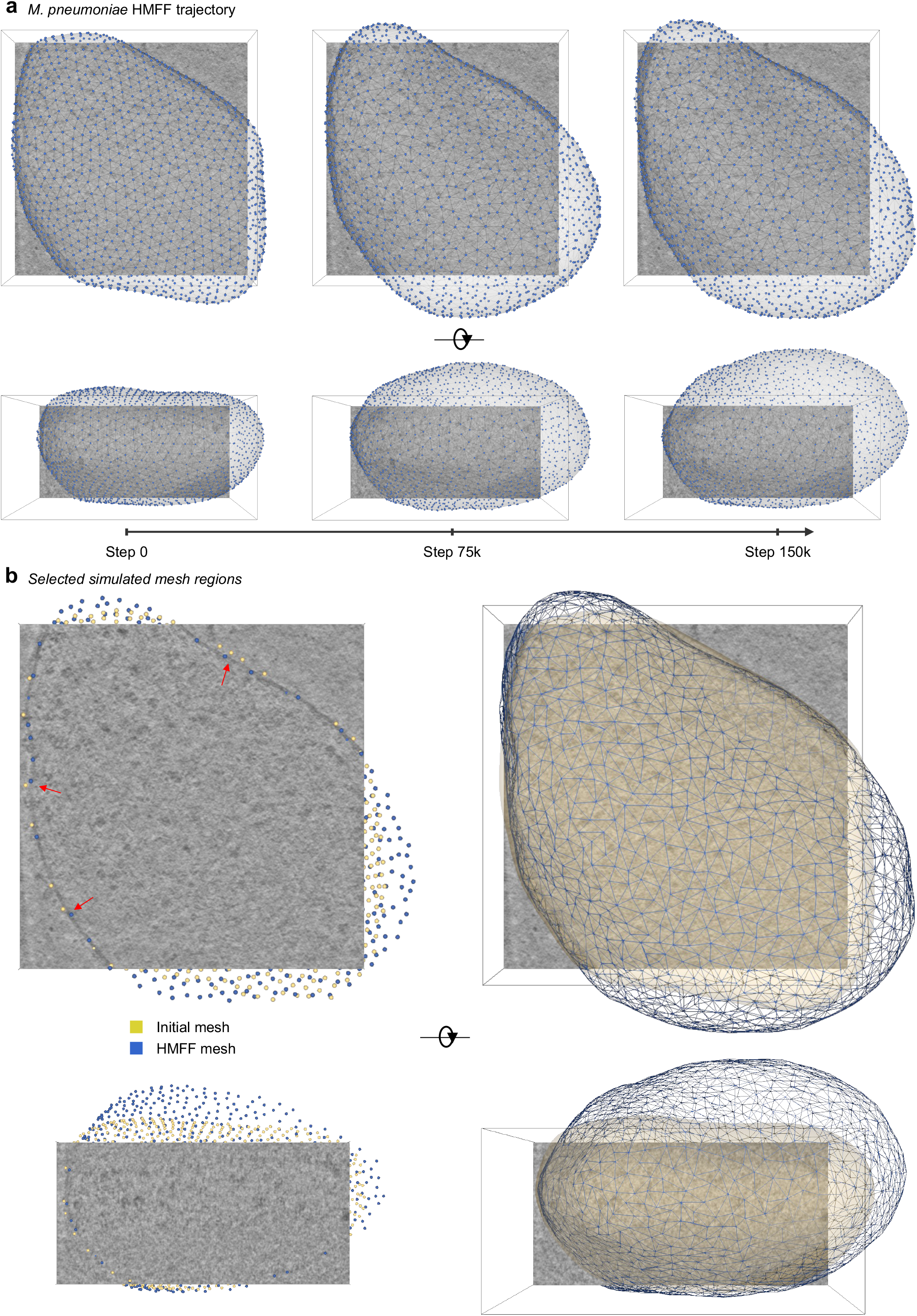

**Fig. S4.**
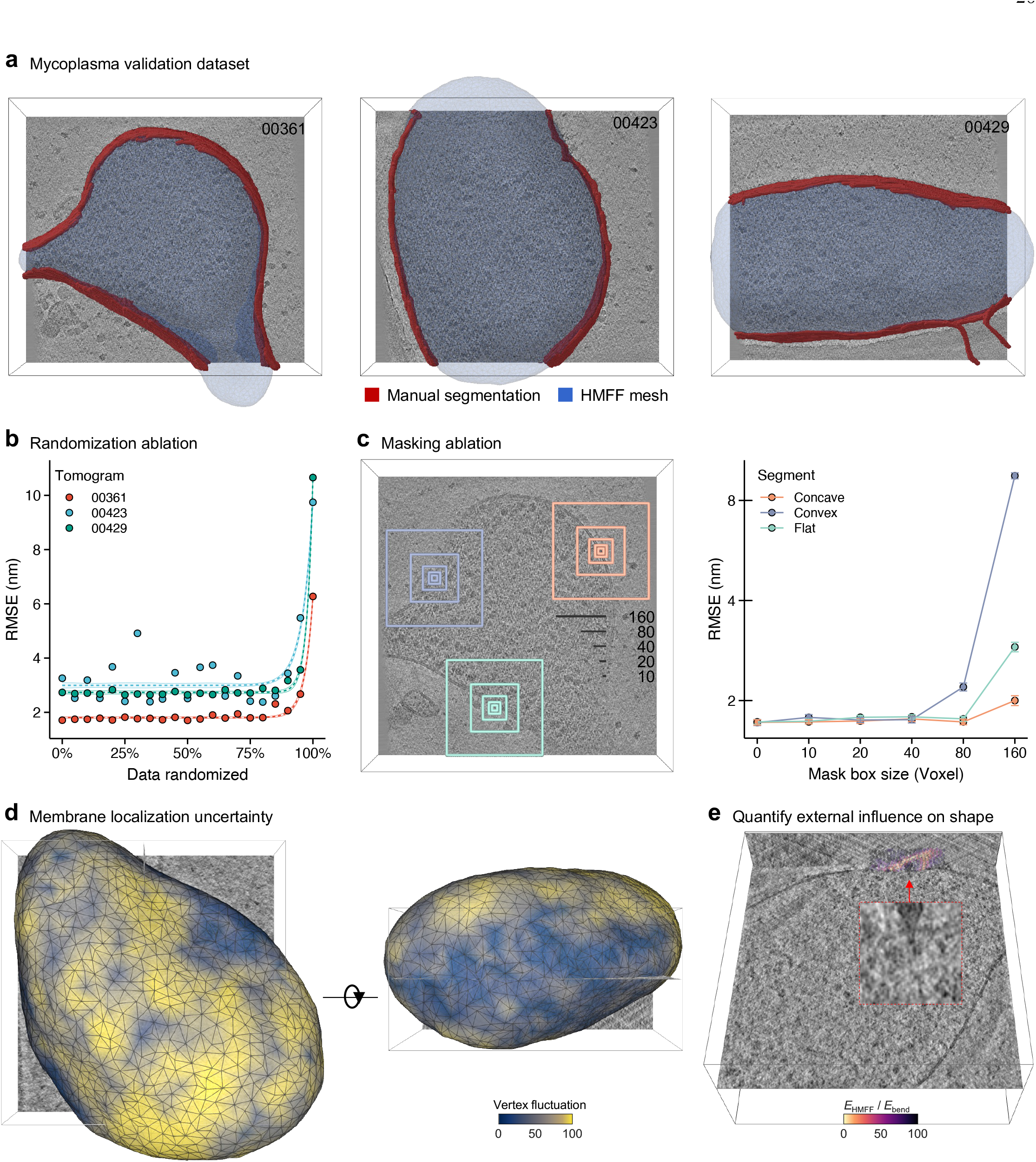
Validation of HMFF on *M. pneumoniae* cryo-ET data. **a**, Three *M. pneumoniae* tomograms with manually segmented membranes and obtained HMFF meshes. **b**, For each tomogram in **a**, increasing fractions of voxel values were replaced with values randomly sampled from the same tomogram, and the RMSE between the manual segmentation and resulting HMFF mesh was measured. Shown is the RMSE over 50k simulation steps, with standard deviation indicated by shaded bands. dashed lines are fits of the form *a* exp(*b x*) + *c*. **c**, Rectangular masks of varying width and fixed *z*-height (80 voxels) were applied at concave, convex, and flat segments of the tomogram (left); RMSE versus mask box size, evaluated as in **b** (right). **d**, Per-vertex fluctuation (standard deviation of position over a 5k step window) on the HMFF mesh of an *M. pneumoniae* cell, shown from two views and colored by quantile. **e**, The particle revealed in the energy decomposition of Fig. 2d, highlighted by a red arrow with magnified inset. The surrounding membrane is overlayed and colored by *E*_HMFF_*/E*_bend_ quantile.

**Fig. S5.**
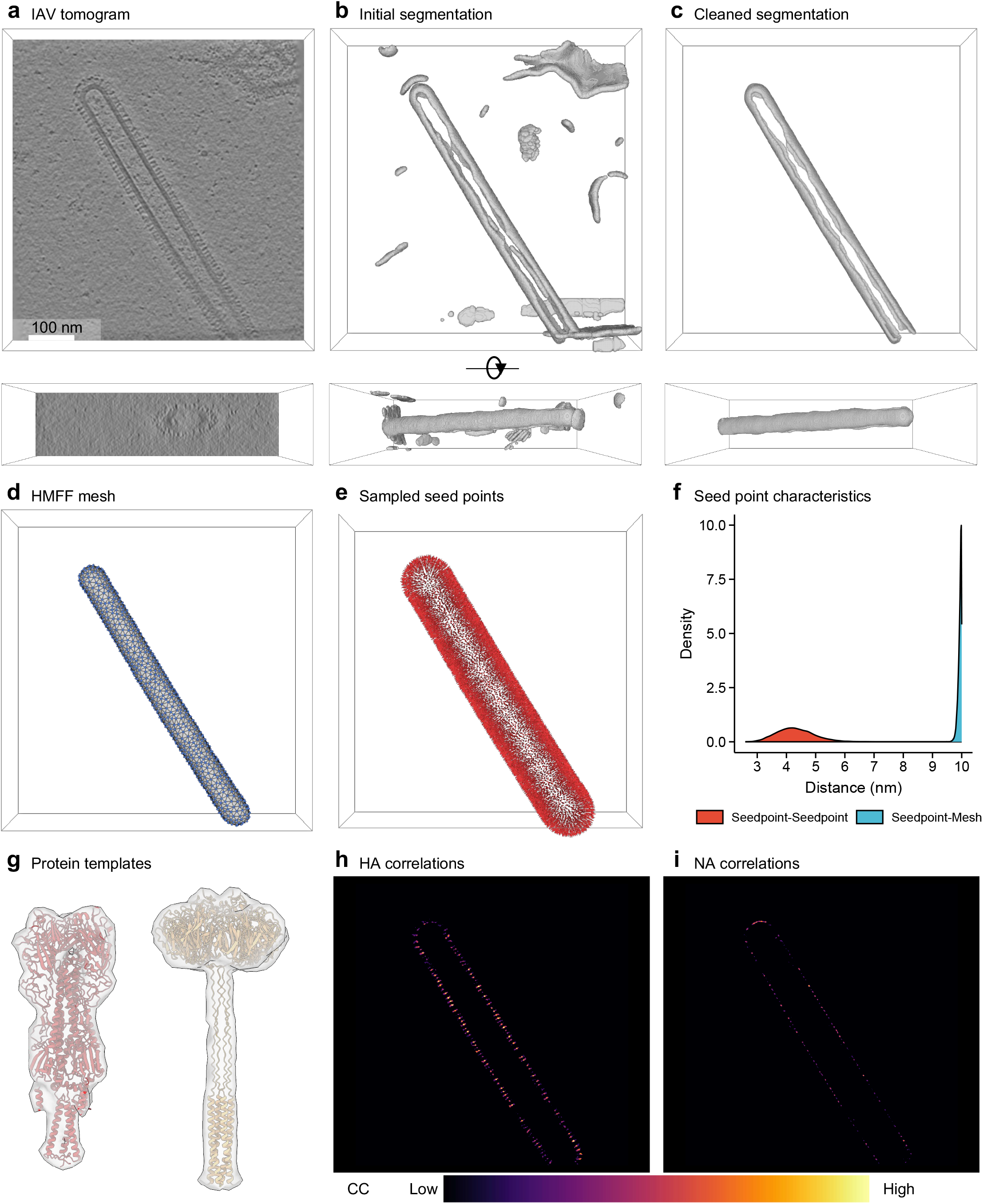
Building a filamentous IAV VLP model. **a**, Band-pass filtered tomographic slice showing a filamentous Influenza A VLP (EMD:11075). **b**, Initial membrane segmentation with visible artifacts. **c**, Membrane segmentation cleaned using Mosaic. **d**, HMFF mesh. **e**, Seed points drawn from the HMFF mesh (**d**) used for template matching. **f**, Distance distribution of seed points, and relative to the HMFF mesh. **g**, HA (red), and NA (orange) structures from Alphafold and corresponding template densities (grey)(see methods IV J). **h**-**i**, Central slice of template matching cross-correlations (CC) for HA (left) and NA (right).

**Fig. S6.**
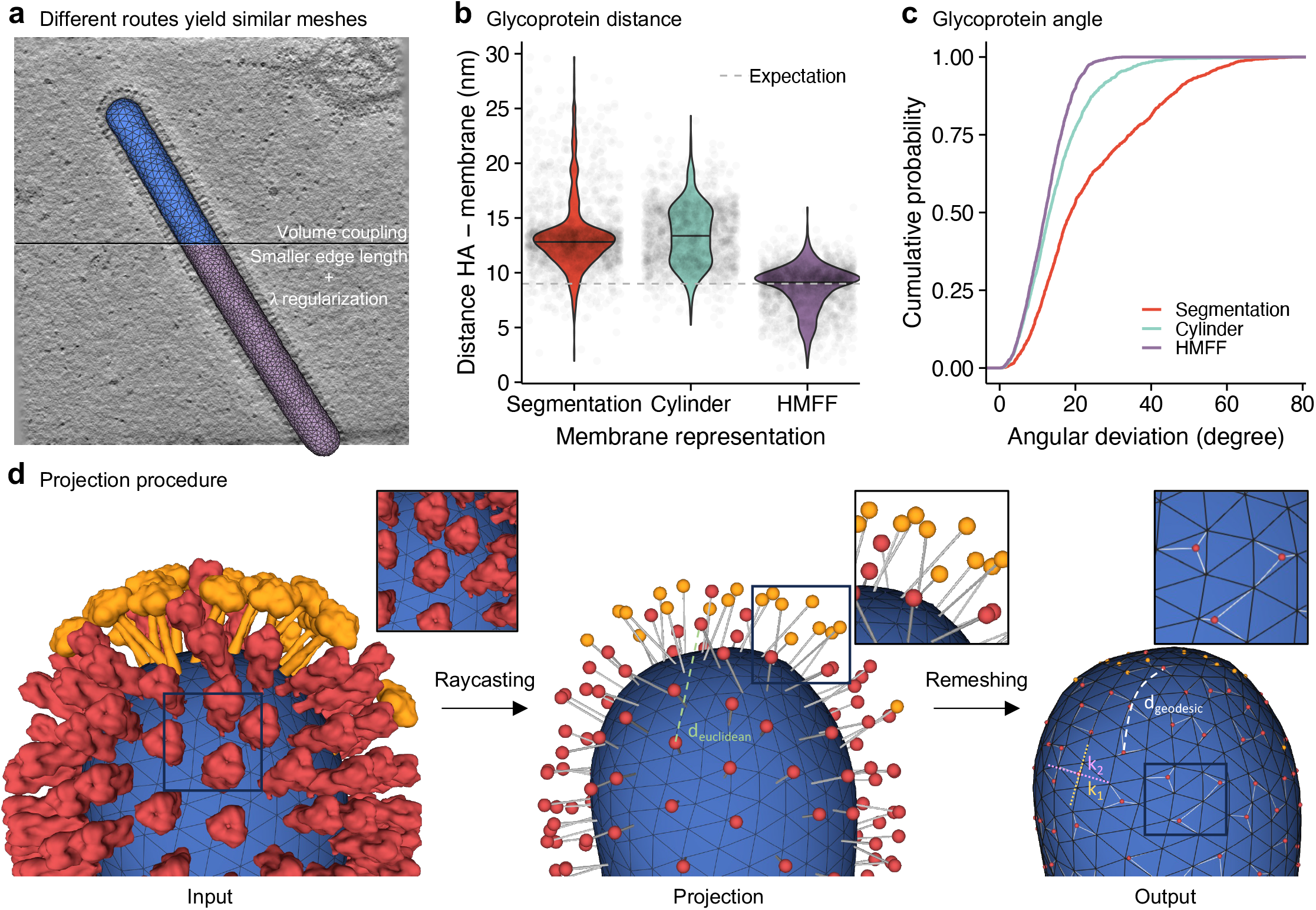
Additional analyses of the IAV VLP HMFF model. **a**, Two HMFF meshes of the same IAV VLP obtained with volume coupling at edge length 11 nm (top), and without volume coupling at reduced edge length 7 nm and *λ* regularization (bottom). Tomogram was band-pass filtered for visualization. **b**, Distance between picked HA proteins and three membrane representations: the cleaned segmentation, a cylinder fit to the segmentation, and the fine-grained *λ*-regularized HMFF mesh from **a**. Dashed line marks the distance between the HA center of mass and the transmembrane region based on the HA structural model. **c**, Cumulative distribution of angular deviation between HA orientations and the local membrane normal for the three representations in **b. d**, Membrane proteins (red/orange) are ray-cast along their normal vectors onto the membrane surface (blue), creating new vertices and edges (white) at intersection points that preserve orientation information from template matching. The extended mesh enables computation of geodesic distances and principal curvatures (*k*_1_, *k*_2_) at precise protein locations.

**Fig. S7.**
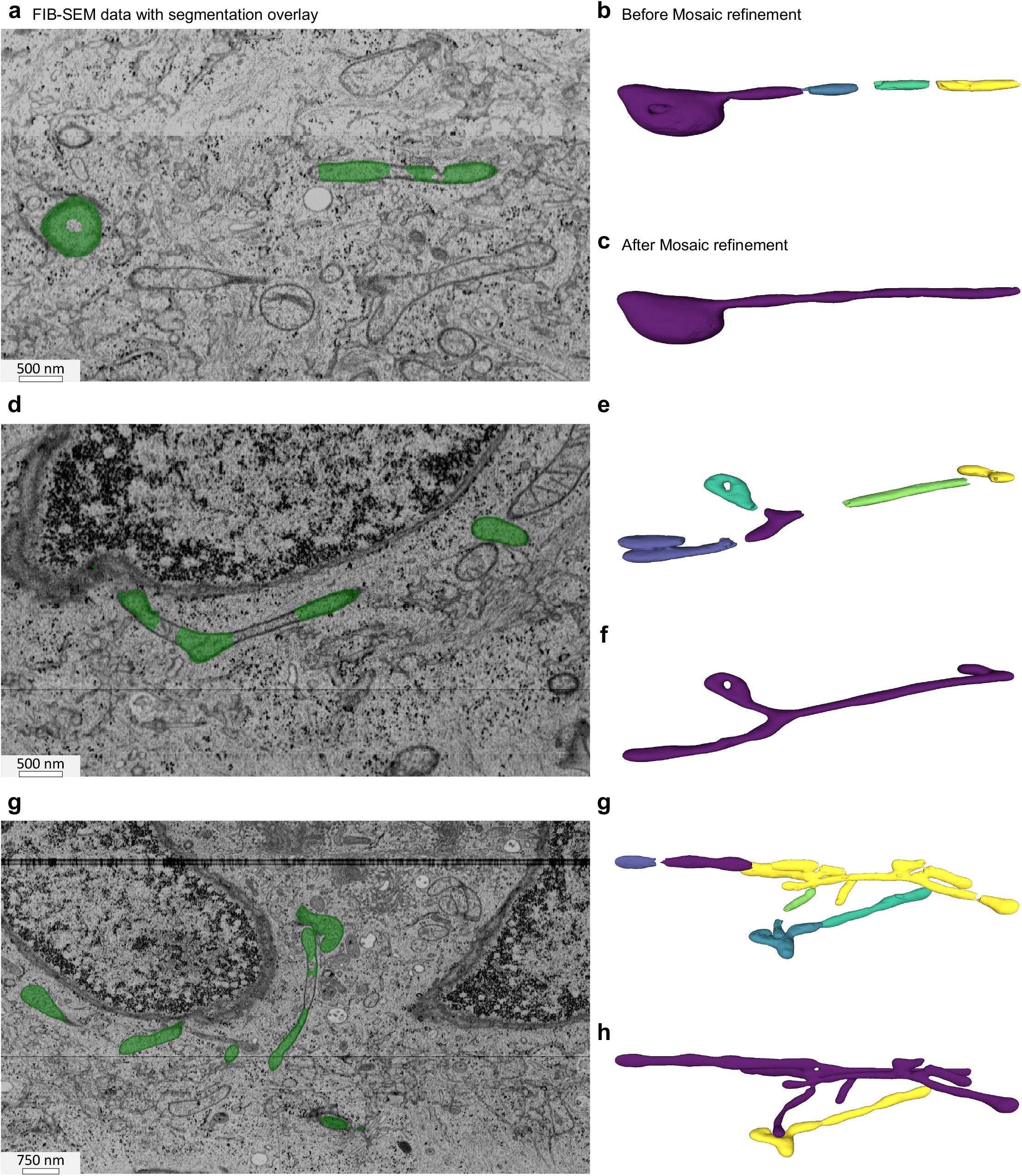
Mitochondrial segmentation quality issues and their resolution using Mosaic. **a, d, g**, Raw FIB-SEM data with original mitochondrial segmentations from Heinrich *et al*. ^22^ overlaid in green. **b, e, h**, Initial meshes obtained from marching cubes applied to the segmentations on the left, with separate entities in the original segmentation shown in different colors. **c, f, i**, Meshes after processing in Mosaic, with the disconnected fragments in (b, e, h) correctly merged into a single connected mitochondrion per row. Rows correspond to distinct mitochondrial morphologies: tubular with bulbous ending (top), branched structure (middle), and complex network with multiple branches (bottom).

**Fig. S8.**
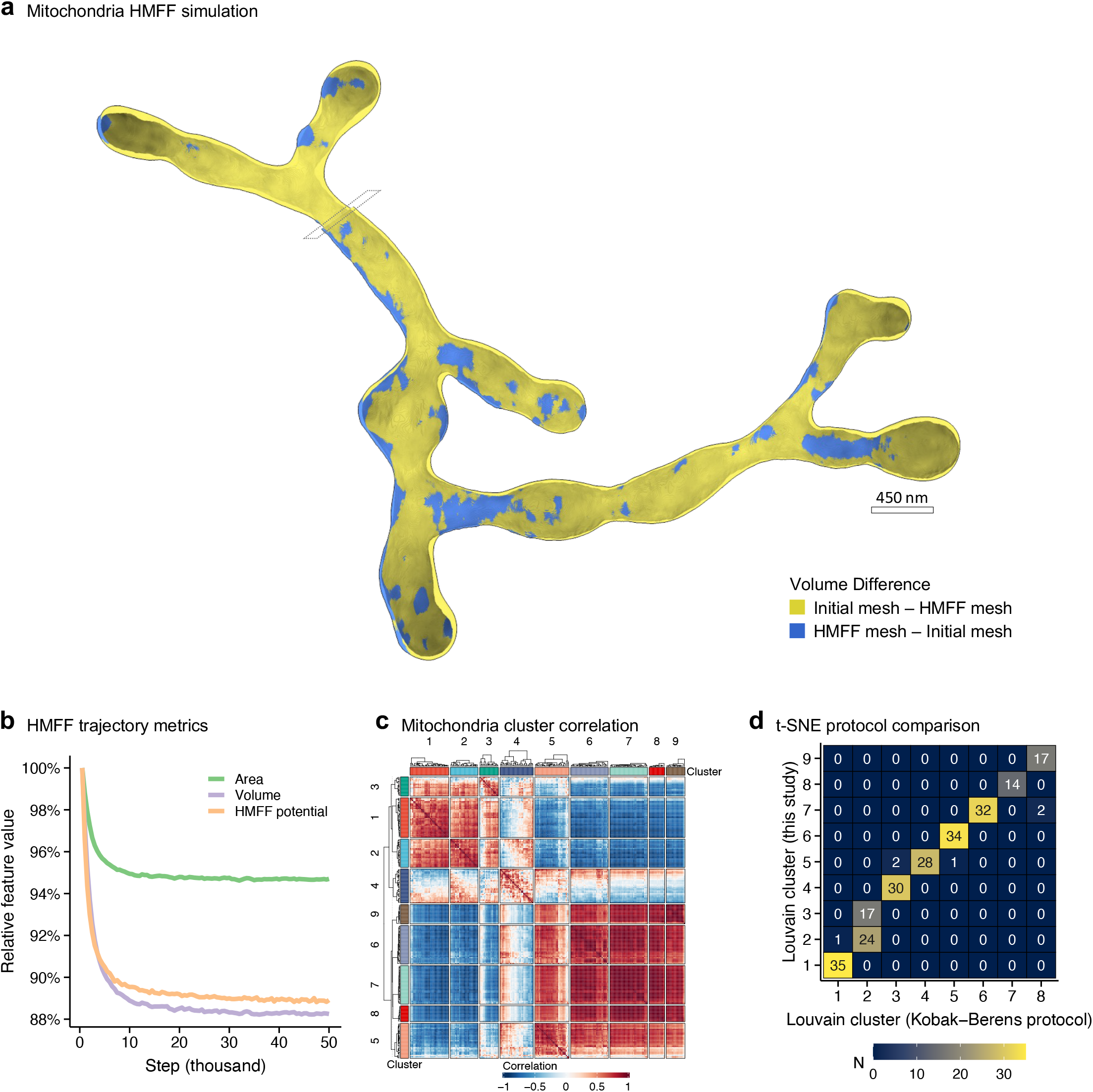
Additional analyses of mitochondrial HMFF simulation and morphological clustering. **a**, Volume difference between the initial and the HMFF mesh for a representative mitochondrion. Yellow marks volume present in the initial mesh but not the HMFF mesh; blue marks the reverse. Dashed lines indicate the image slice plane. **b**, HMFF simulation trajectory: HMFF energy, surface area, and volume over simulation steps, each max-scaled for comparison. **c**, Hierarchical clustering of mitochondrial morphology descriptors with Pearson correlation heatmap. **d**, Comparison of Louvain cluster assignment obtained from the t-SNE embedding in Fig. 5d, and an alternative embedding using the protocol introduced by Kobak and Berens ^100^ (see methods IV N). Numbers indicate number of mitochondria within a cluster.

**Fig. S9.**
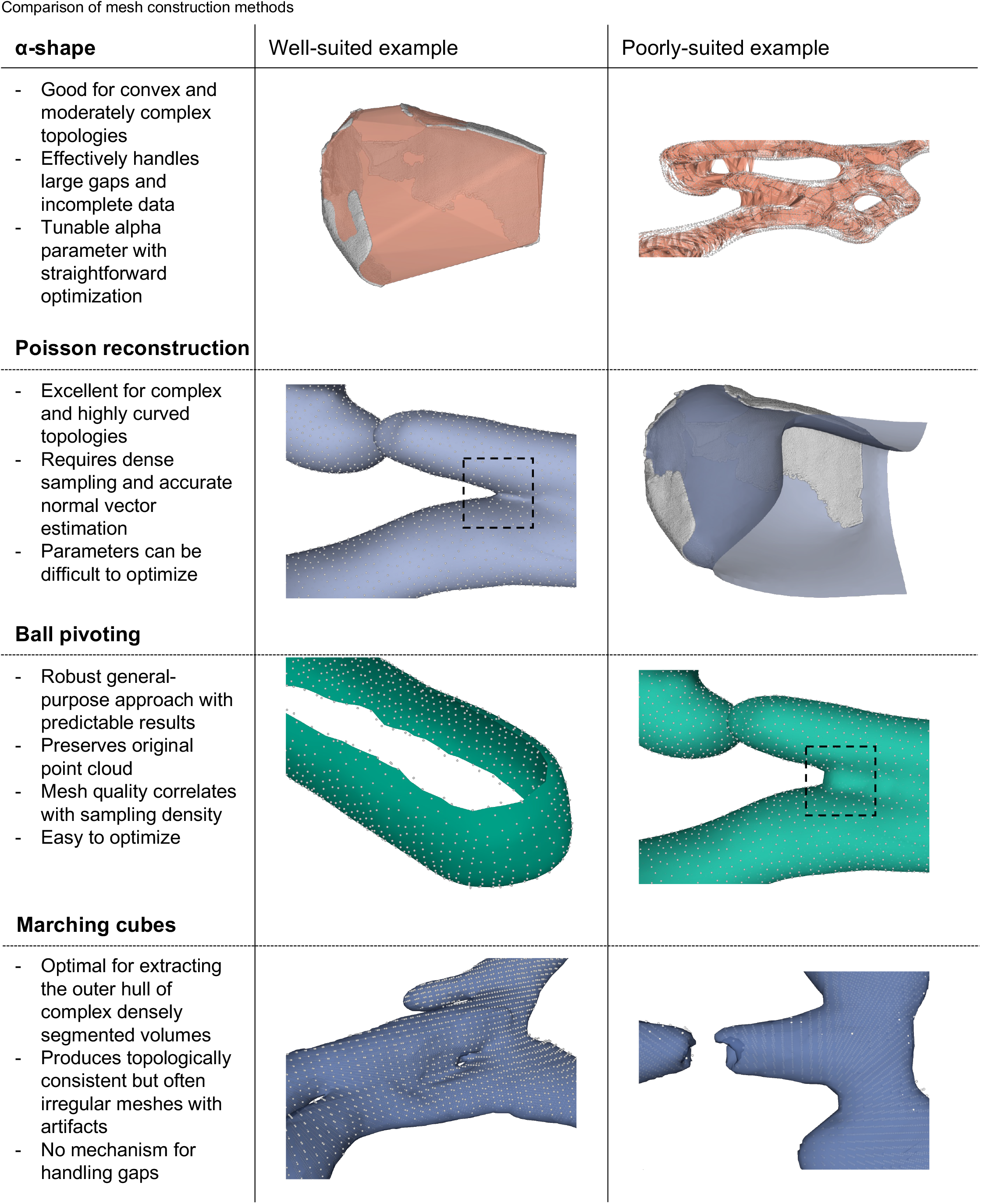
Comparison of key mesh construction methods implemented in Mosaic. Illustrated are meshing approaches, showing well-suited examples where a method excels and poorly-suited examples where it faces limitations. Points shown in grey indicate the input used for triangulation. *α*-shape, Poisson reconstruction, and Ball pivoting operate on point clouds, while marching cubes processes volumetric data.

**Fig. S10.**
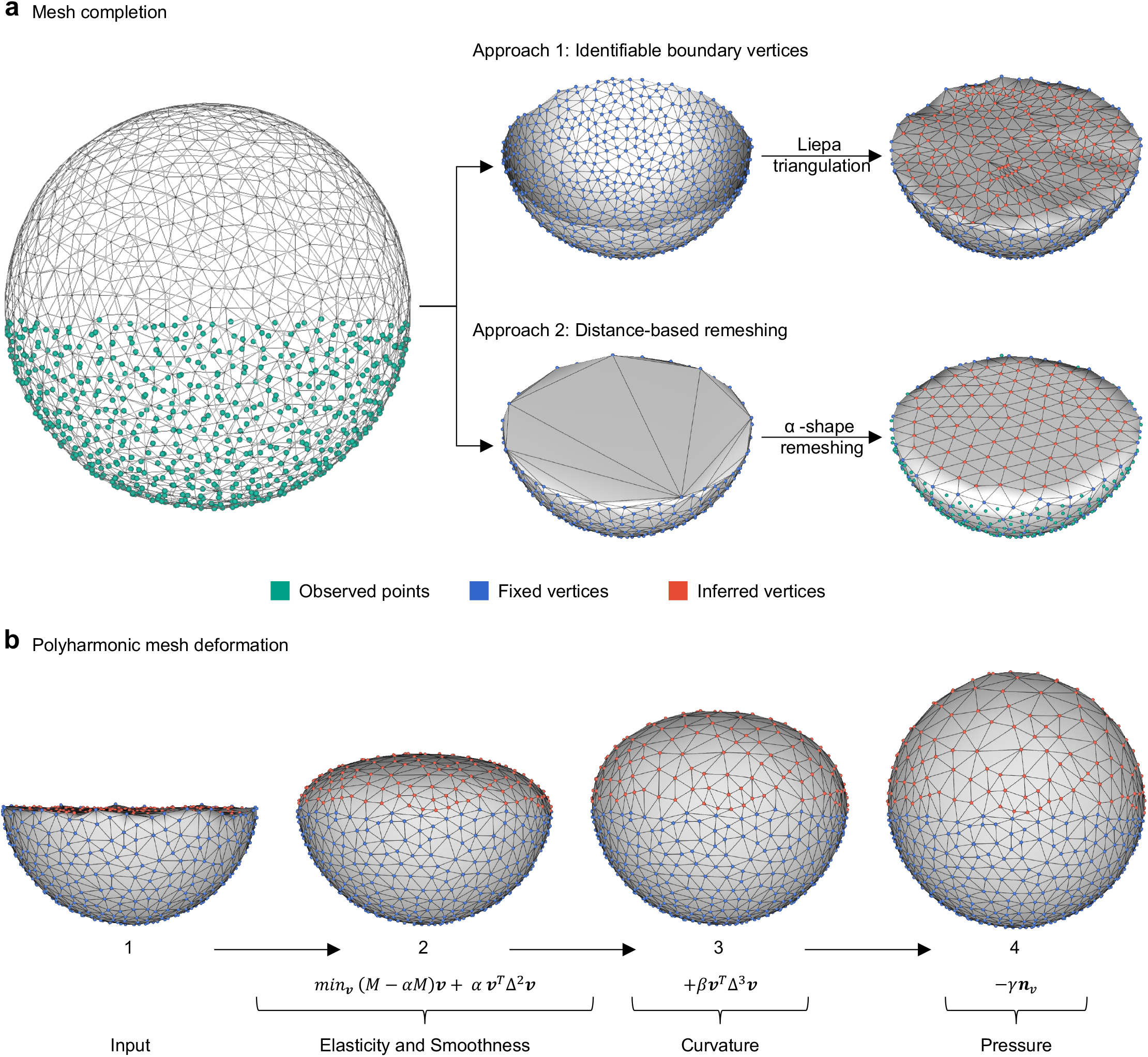
Mesh completion strategies for generating biological membrane models. **a**, Two complementary approaches for completing partial membrane structures: when boundary vertices are identifiable (top), Liepa triangulation preserves local geometry while filling missing regions; for disconnected structures (bottom), distance-based remeshing with *α*-shapes reconstructs closed surfaces. In both approaches, triangulations are based on observed points (green), with fixed vertices (blue) originating from the observation and inferred vertices (red) from the completion strategy. **b**, Progressive deformation of inferred vertices guided by biophysical principles. Inferred vertices (red) undergo optimization to assimilate into the geometry dictated by fixed vertices (blue). This procedure integrates elasticity, smoothness, curvature, and pressure terms (labeled 2-4, see methods IV D 2), ensuring the completed regions seamlessly connect with observed structures while maintaining properties required for HMFF simulation.

Mov. M1. HMFF simulations of a planar membrane in a Gaussian density field with different coupling constants (*ξ*). As *ξ* increases from 0 (no coupling) to 10 (strong coupling), the membrane progressively shows stronger and more rapid adaptation to the density.

Mov. M2. *M. pneumoniae* membrane segmentation process. Comparative visualization of raw cryo-ET data (left), initial deep-learning segmentation (middle), and manually refined segmentation after processing in Mosaic (right).

Mov. M3. *M. pneumoniae* HMFF simulation. The membrane mesh evolves during simulation, balancing experimental observation with physical membrane properties. A frame corresponds to 1,000 simulation steps.

Mov. M4. HMFF simulation of *M. pneumoniae* without boundary padding, showing mesh erroneous flattening and attachment to tomogram boundaries.

Mov. M5. IAV VLP membrane segmentation. Comparative visualization of raw cryo-ET data (left), initial deep-learning segmentation (middle), and manually refined segmentation after processing in Mosaic (right).

Mov. M6. IAV VLP HMFF simulation. The membrane mesh evolves during the simulation, moving out of the M1 protein layer and stabilizing at the experimentally observed membrane. A frame corresponds to 1,000 simulation steps.

Mov. M7. IAV VLP model with glycoprotein distribution. HMFF mesh showing the distinct spatial organization of hemagglutinin (HA) and neuraminidase (NA) proteins across the viral surface. This visualization demonstrates how the HMFF mesh can be used as a framework for studying membrane proteins.

Mov. M8. Mitochondria HMFF simulation. The mesh adapts to FIB-SEM density data while maintaining overall membrane geometry. Each frame corresponds to 500 simulation steps.

Mov. M9. Three-dimensional rendering of Mitochondria HMFF mesh showcasing the complex branched morphology present in the experimental data.

Mov. M10. HeLa cell ultrastructure mesh reconstruction. Integrated view of cellular compartments including nucleus (orange), endoplasmic reticulum (purple), Golgi apparatus (blue), and mitochondria (green), demonstrating cellular-scale membrane modeling capabilities at the micrometer scale.

